# Efficient and accurate detection of viral sequences at single-cell resolution reveals putative novel viruses perturbing host gene expression

**DOI:** 10.1101/2023.12.11.571168

**Authors:** Laura Luebbert, Delaney K. Sullivan, Maria Carilli, Kristján Eldjárn Hjörleifsson, Alexander Viloria Winnett, Tara Chari, Lior Pachter

## Abstract

There are an estimated 300,000 mammalian viruses from which infectious diseases in humans may arise. They inhabit human tissues such as the lungs, blood, and brain and often remain undetected. Efficient and accurate detection of viral infection is vital to understanding its impact on human health and to make accurate predictions to limit adverse effects, such as future epidemics. The increasing use of high-throughput sequencing methods in research, agriculture, and healthcare provides an opportunity for the cost-effective surveillance of viral diversity and investigation of virus-disease correlation. However, existing methods for identifying viruses in sequencing data rely on and are limited to reference genomes or cannot retain single-cell resolution through cell barcode tracking. We introduce a method that accurately and rapidly detects viral sequences in bulk and single-cell transcriptomics data based on highly conserved amino acid domains, which enables the detection of RNA viruses covering over 100,000 virus species. The analysis of viral presence and host gene expression in parallel at single-cell resolution allows for the characterization of host viromes and the identification of viral tropism and host responses. We applied our method to identify putative novel viruses in rhesus macaque PBMC data that display cell type specificity and whose presence correlates with altered host gene expression.

## Introduction

There are an estimated 10^31^ virions on Earth, which amounts to 10 million virions for every star in the known universe^1,2^. Viruses inhabit oceans, forests, deserts, and human tissues such as the lungs, blood, and brain. There are an estimated 300,000 mammalian viruses^3^ from which infectious diseases in humans may arise^4^. However, only 261 species have been detected in humans^5^. Many of these have been implicated in complex diseases such as heart disease and cancer. Recent studies suggest that viruses also play a major, unexpected role in common neurodegenerative disorders such as Alzheimer’s, Parkinson’s, and multiple sclerosis^6-8^. Accurate detection of viral infections is crucial to understanding viral impact on human health.

Of the 261 known disease-causing viruses, 206 fall into the realm of *Riboviria*^5^, which includes all RNA-dependent RNA polymerase (RdRP)-encoding RNA viruses and RNA-dependent DNA polymerase (RdDP)-encoding retroviruses. Amongst many others, these include Corona-, Dengue, Ebola-, Hepatitis B, influenza, Measles, Mumps, Polio-, West Nile, and Zika viruses. Most existing workflows for detecting viruses in transcriptomics data rely on the availability of pre-assembled reference genomes. Currently, NCBI RefSeq hosts 8,694 *Riboviria* reference genomes—a diminutive fraction of *Riboviria* viruses. Pioneering work by Edgar *et al.*^9^ leveraged a well-conserved amino acid sub-sequence of the RdRP, called the ‘palmprint’, to identify RNA viruses in 5.7 million globally and ecologically diverse sequencing samples from the Sequence Read Archive (SRA). Their method’s independence from pre-computed indices allowed alignment to diverged sequences, leading to the discovery of thousands of novel viruses. This effort resulted in a consensus of 296,623 unique RdRP-containing amino acid sequences, henceforth referred to as ‘PalmDB’. Clustering palmprints into species-like operational taxonomic units (sOTUs) yielded 146,973 known as well as novel sOTUs^9^. Compared to the 8,694 *Riboviria* reference genomes currently available on NCBI, this translates to a more than 16x increase in the number of viruses that can be detected. The actual number of virus species that can be detected using the PalmDB is likely even higher due to RdRP sequence conservation across *Riboviria* (Extended Data Fig. 1). sOTUs serve to approximate taxonomic assignment^9,10^ and allow species-level virus identification for 40,392 sequences in the PalmDB.

The increasing use of high-throughput next-generation sequencing (NGS) methods in molecular biology research, agriculture, and healthcare provides an opportunity for the cost-effective surveillance of viral diversity and the investigation of virus-disease correlations^11,12^. Specifically, single-cell genomics technologies make possible, in principle, the characterization of viruses at single-cell resolution. We expanded the RNA sequencing data preprocessing tool kallisto^13^ to support the detection of viral RNA using the amino acid database PalmDB. To our knowledge, this is the only existing method capable of translated alignment while retaining single-cell resolution. The small size of PalmDB (36 MB) enables efficient detection of orders of magnitude more viruses than detection based on (NCBI) reference genomes. Moreover, operating in the amino acid space yields a method robust to silent nucleotide mutations.

## Results

### Translated alignment of nucleotide sequences to an amino acid reference with kallisto enables efficient, accurate detection of RNA viruses at single-cell resolution

Most existing methods to detect viral sequences either (i) rely on and are limited to (NCBI) reference genomes^14–23^, (ii) are not able to perform translated alignment of nucleotide data to an amino acid reference^24–26^, or (iii) are unable to retain single-cell resolution through cell barcode tracking^27,28^ (Fig. 2b). We expanded the bulk and single-cell RNA-seq preprocessing tool kallisto^13^ to allow translated search and validated its use in combination with PalmDB for the detection of virus-like sequences in single-cell and bulk RNA sequencing data. PalmDB is a database of 296,623 unique RdRP-containing amino acid sequences, representing 146,973 virus species^9^. Fig. 2a provides an overview of the number of entries per taxonomy in the NCBI and PalmDB databases. The figure can also be viewed interactively here: tinyurl.com/4dzwz5ny.

The translated alignment is performed by first reverse translating the amino acid reference sequences and all possible reading frames (three forward and three reverse) of the nucleotide sequencing reads to comma-free code (Fig. 1)^29^. A comma-free code is a set of k-letter ‘words’ selected such that any off-frame k-mers formed by adjacent letters do not constitute a ‘word’, and will thus be interpreted as ‘nonsense’. For k=3 (a triplet code) and 4 letters (e.g., ‘A’, ‘T’, ‘C’, and ‘G’), this results in exactly 20 possible words (theorem shown in Fig. 1), which equals the number of amino acids specified by the universal genetic code. Due to the serendipity of these numbers, Crick *et al.* hypothesized the genetic code to be a comma-free code in 1957^30^. The impossibility of off-frame matches makes comma-free codes highly appropriate for translated alignment (Extended Data Fig. 10, Methods). The *de Bruijn* graph generated from the reverse translated PalmDB sequences groups viruses of the same taxonomies, indicating that within-taxonomy similarity is conserved in comma-free space as expected (Extended Data Fig. 4d). Finally, the six reading frames of the sequencing reads translated to comma-free code are pseudoaligned to the reference sequences reverse translated to comma-free code. If several reading frames of the same read produce alignments, the best frame is chosen (Fig. 1, Methods).

**Fig. 1:**
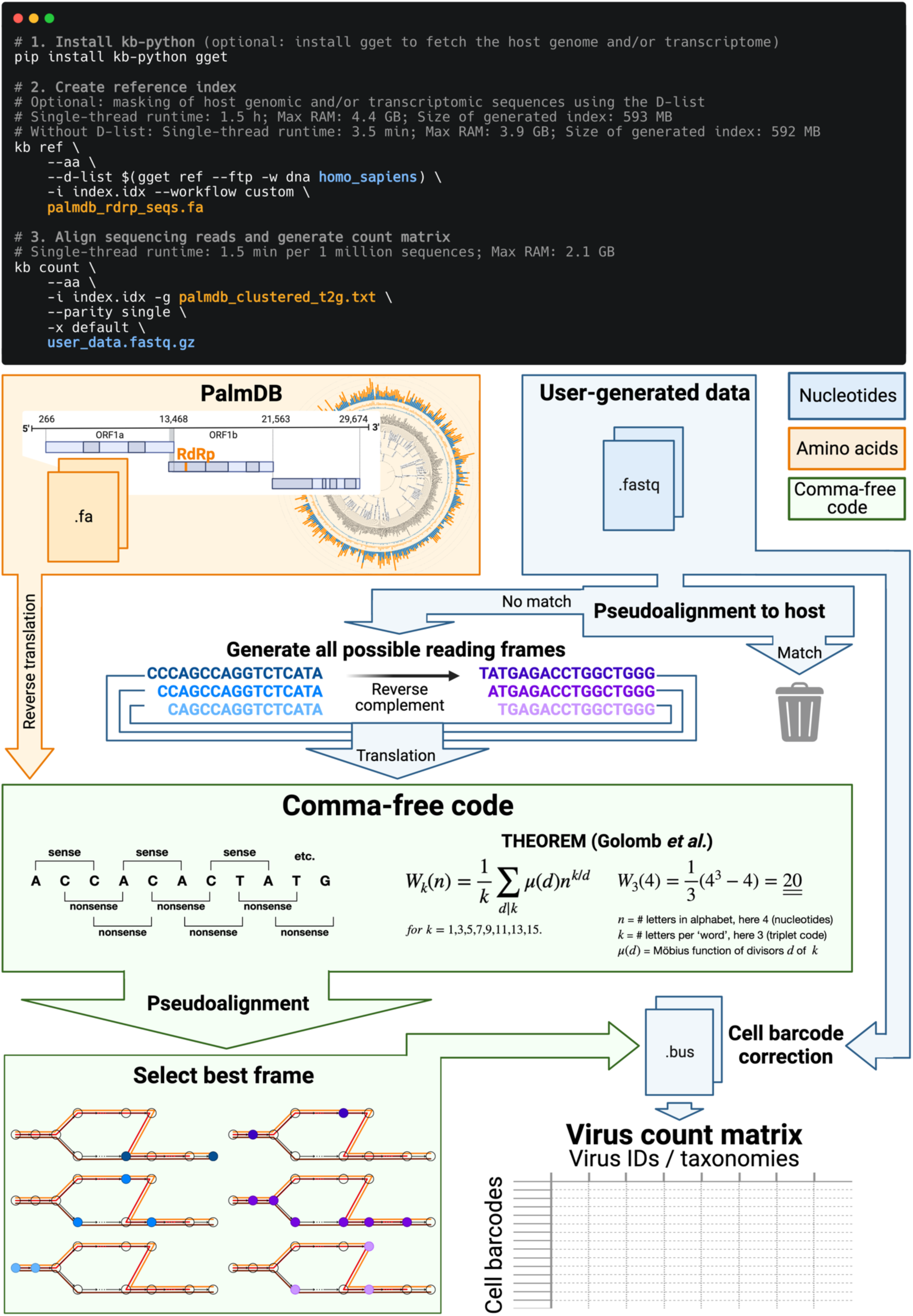
Schematic overview of the kallisto translated search front- and back-end. The front-end is similar to kallisto-bustools workflows as previously described^43,44^. The user provides sequencing data, usually in the form of FASTQ files, as well as a reference FASTA file containing amino acid sequences to align the nucleotide sequencing data against. The novel argument ‘--aa’ activates the translated search alignment. In the example shown here, the reference file consists of the PalmDB amino acid RdRP sequences contained in ‘palmdb_rdrp_seqs.fa.’ The ‘palmdb_clustered_t2g.txt’ file groups virus IDs with the same taxonomy across all main taxonomic ranks like transcripts of the same gene (see Methods). Both files are available here: https://tinyurl.com/4wd33rey. During the generation of the reference index with ‘kb ref’, the D-list option may be used to mask host genomic and/or transcriptomic sequences, as further discussed in this manuscript. Here, human genomic sequences fetched from Ensembl using gget^45^ are masked using the D-list. The reference index only needs to be generated once, and precomputed PalmDB reference indices for human and mouse hosts are available here: https://tinyurl.com/aaxyy8v8. Following the generation of a reference index, the sequencing reads are pseudoaligned to the reference index, and a count matrix is generated using the ‘kb count’ command. The ‘-x’ argument is used to define the sequencing technology. In the example code, the minimum required user input is marked in orange (amino acid space) and blue (nucleotide space). In the kallisto translated search back- end, the reference amino acid sequences and the nucleotide sequencing reads are translated into a non-redundant comma-free code. For the nucleotide sequences, the translation occurs in all six possible reading frames (three forward and three reverse frames). The pseudoalignment is performed in the comma-free code space and is compatible with the kallisto cell barcode tracking which enables analysis at single-cell resolution. The workflow generates a cell barcode by virus ID count matrix.

The workflow can be executed in three lines of code, and computational requirements do not exceed those of a standard laptop (Fig. 1). Building on kallisto’s versatility, the workflow is compatible with all state-of-the-art single-cell and bulk RNA sequencing methods, including but not limited to 10x Genomics, Drop-Seq^31^, SMART-Seq^32^, SPLiT-Seq^33^ (including Parse Biosciences), and spatial methods such as Visium.

Validation testing was performed using different bulk and single-cell RNA sequencing datasets with known infections with severe acute respiratory syndrome coronavirus 2 (SARS-CoV-2) or *Zaire ebolavirus* (ZEBOV)^34–38^. In these tests, translated search with kallisto and PalmDB was able to detect the viral RNA and correctly assign species-level taxonomy at counts correlating with viral loads measured by RT-qPCR or RNA-ISH, regardless of the technology used to generate the data (Fig. 3a). Fig. 3b provides an overview of the robustness of the taxonomic assignment across all available taxonomic ranks after reverse translated RdRP sequences were aligned to the PalmDB with kallisto translated search. At the species level, 96.76 % of 296,561 sequences were assigned the correct taxonomy, 0.007 % were assigned an incorrect taxonomy, 0.37 % could not be unambiguously matched to a single virus (they were multimapped), and 2.86 % were not aligned. This confirms that the sequence transformation introduced by the kallisto translated search pipeline retains taxonomic assignments with up to species-level specificity.

Next, we sought to confirm that kallisto translated search with PalmDB correctly identifies sequences that originate from the RdRP gene. To this end, we selected a subset of 100,000,000 reads obtained using Seq-Well sequencing of macaque peripheral blood mononuclear cell (PBMC) samples obtained at 8 days post-infection with ZEBOV^37^ (see Methods). We aligned the reads to the PalmDB amino acid sequences with kallisto translated search. We also aligned the reads to the complete ZEBOV nucleotide genome using Kraken2 (standard nucleotide alignment)^27^. Aligned reads from both alignments were extracted and realigned to the ZEBOV genome using bowtie2^39^, a BAM file was created using SAMtools^40^ and the alignment was subsequently visualized NCBI Genome Workbench^41^. The visualized alignments are shown in Extended Data Fig. 2 and confirmed that kallisto translated search accurately and comprehensively detected ZEBOV RdRP sequences.

We then tested whether our translated search method is robust to single nucleotide mutations, which occur at a relatively high rate in RNA viruses of up to 10^−4^ substitutions per nucleotide site per cell infection^42^. We added random single nucleotide base substitutions to 676 ZEBOV nucleotide RdRP sequences identified during the alignment described in the previous paragraph^37^, then assessed the frequency of correct taxonomic classification (recall percentage) by kallisto translated search, in comparison to the current state-of-the-art translated search tool, Kraken2 (translated search). kallisto translated search correctly recalled up to 27.5 % more viral RdRP sequences than Kraken2 (translated search) (Fig. 2c). Moreover, kallisto translated search was more robust than aligning to the complete nucleotide genome with the standard kallisto workflow at mutation rates > 4 % (Fig. 2c), which emphasizes the advantage of operating in the amino acid space. While the Kraken2 (translated search) and the kallisto standard workflow were given only the correct virus as a reference (here, ZEBOV), kallisto translated search had to distinguish between all viruses contained in the PalmDB and identify the correct taxonomy. kallisto translated search was able to maintain > 90 % precision in the species-level taxonomic assignment at mutation rates up to 12 % (Extended Data Fig. 4b).

**Fig. 2:**
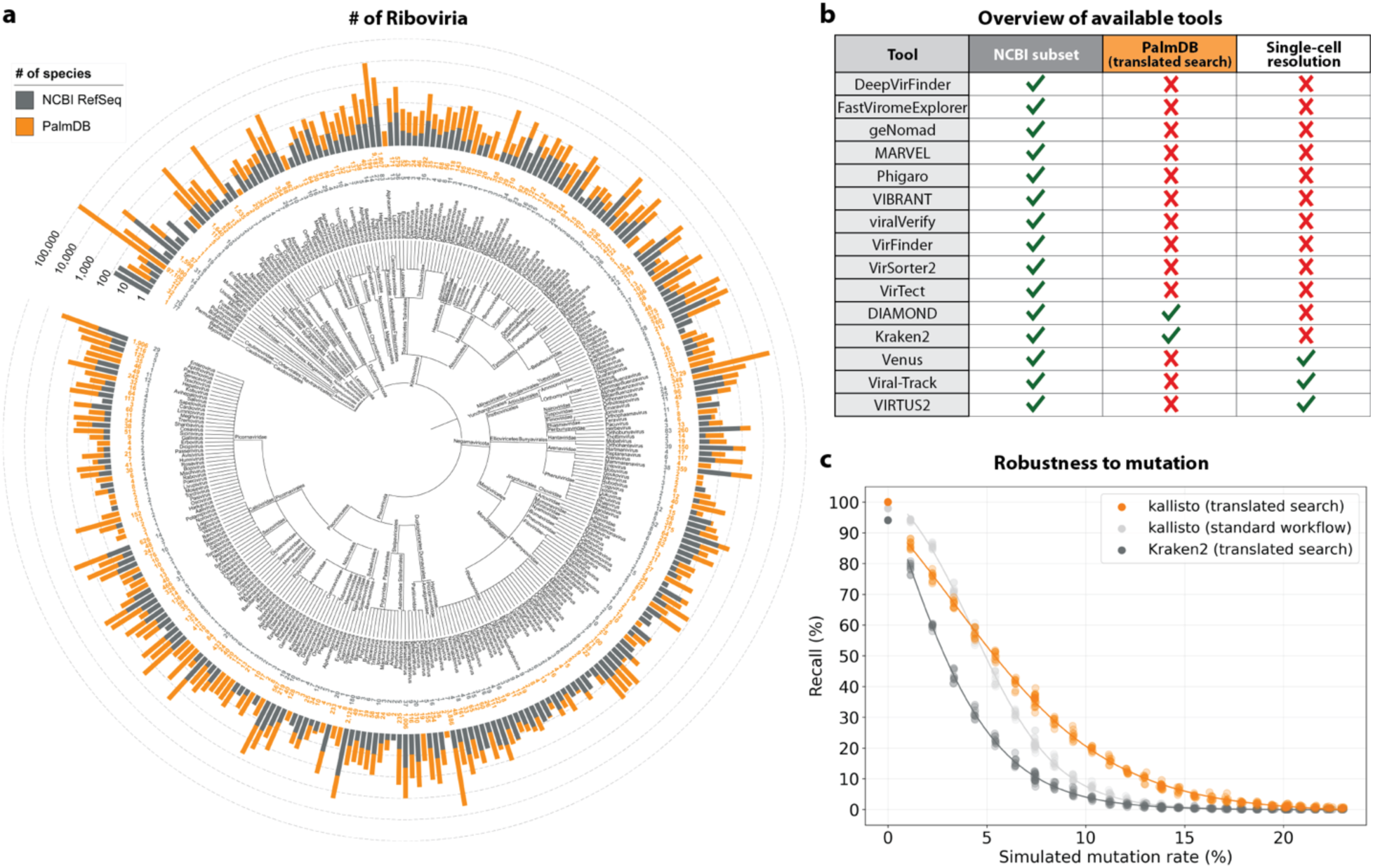
**a**, Phylogenetic tree of the taxonomies of viral sequences/genomes included in the PalmDB sOTUs and NCBI RefSeq databases from phylum to genus. Barplots indicate the number of sequences/species available for each taxonomy in each database. The tree was generated with iTOL^46^. This plot can also be viewed interactively here: <u>tinyurl.com/4dzwz5ny</u>. **b**, Overview of available tools for the detection of viral sequences in next-generation sequencing data^14–26,28,47^, and their ability to align to NCBI RefSeq nucleotide genomes, perform translated alignment of nucleotide data against an amino acid reference, and retain single-cell resolution through cell barcode tracking. **c**, Mutation-Simulator^48^ was used to add random single nucleotide base substitutions to 676 ZEBOV RdRP sequences obtained by Seq-Well sequencing^37^ at increasing mutation rates. We performed 10 simulations per mutation rate. The sequences were subsequently aligned using kallisto translated search against the complete PalmDB, Kraken2 translated search against the RdRP amino acid sequence of ZEBOV with a manually adjusted NCBI Taxonomy ID to allow compatibility with Kraken2, and kallisto standard workflow against the complete ZEBOV nucleotide genome (GCA_000848505.1). The plot shows the recall percentage of the 676 sequences for each of the 10 simulations at each mutation rate. Each was fitted with an inverse sigmoid for mutation rates > 0.

We next sought to investigate whether viral species not included as species-like operational taxonomic units (sOTUs) in the reference PalmDB database could be detected based on the conservation of the RdRP gene. To do this, we removed all *Ebolavirus* species, all *Ebolavirus* genera, and all members of the *Filoviridae* family from the reference, and subsequently aligned the 676 ZEBOV RdRP sequences obtained by Seq-Well sequencing^37^. In each scenario, a subset of sequences aligned to the nearest remaining relative based on the main taxonomic rank (Extended Data Fig. 1). This suggests that kallisto translated search can detect the highly conserved RdRP of a large number of viral species, beyond the number of sequences in the PalmDB database, while still providing reliable sOTU-based taxonomic assignment of lower-rank taxonomies.

### Read and virus filtering

A common problem that arises during the identification of microbial sequences is the cross-species contamination of reference genome databases, such as the ubiquitous contamination of bacterial genomes with human DNA^49–51^. The PalmDB is not a curated database, and it is possible that some virus-like sequences in the PalmDB are not derived from viruses. This can lead to the misclassification of host reads as bacterial or viral, suggesting the presence of microbes that were not truly present. The misclassification of host reads as viral can be prevented by removing host reads prior to the alignment to the viral reference. However, conservatively removing host reads will also remove sequences of endogenous viral elements, which are very abundant in vertebrate genomes^52^ and may lead to the removal of viral sequences that were truly present. Hence, there are two goals: (i) removing host reads to prevent the misclassification of host reads as viral while (ii) comprehensively identifying the virome within a sample.

In some instances, it is impossible to unambiguously determine whether a read originated from the host or a virus/microbe during the alignment. For example, an analysis of cancer microbiomes by Poore *et al.*^55^ identified the presence of several bacterial genera. While the reanalysis of this data by Gihawi *et al.*^49^ using highly conservative host sequence masking led to a significant decrease in the number of reads identified for each bacterial genus, many bacterial genera remained identified. We found that conservative host masking can lead to the removal of sequences originating from a confirmed viral infection, as was the case for ZEBOV here (Fig. 4c and 5a). Hence, it is unclear whether removing all ambiguous reads prior to downstream analysis is correct or results in biologically relevant data being thrown out. Thus, we developed easy-to-use, flexible, and efficient sequence masking methods, which allow the identification of reads that align to both host and virus/microbe while also preserving these reads for downstream analysis. We used these masking methods in combination with PalmDB for the identification of viral sequences. However, they can be applied to mask any nucleotide or amino acid reference and are able to retain single-cell resolution.

**Fig. 3:**
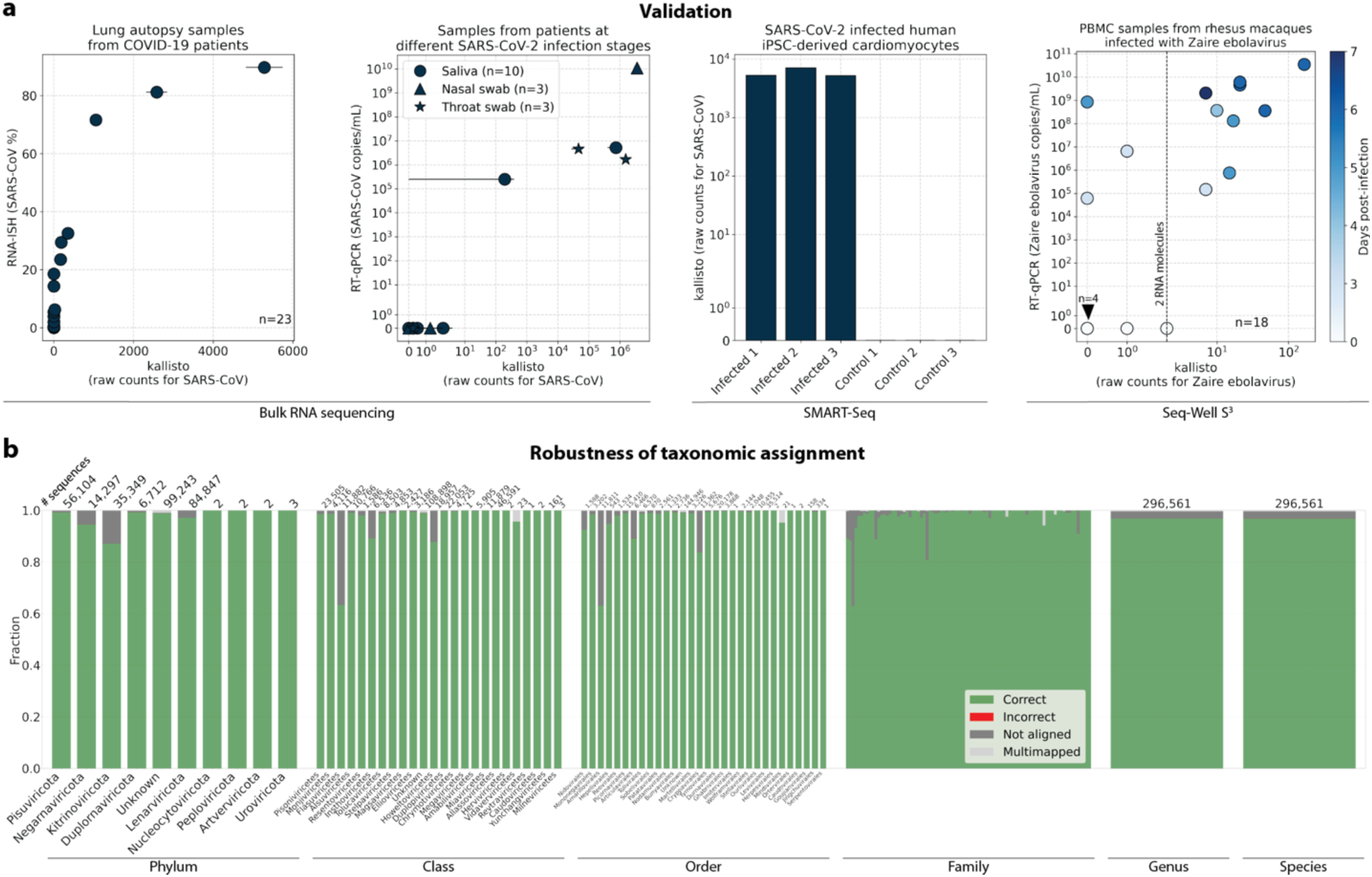
**a**, Sequencing data from samples with a known viral infection and sequenced using different bulk and single-cell RNA sequencing technologies was aligned to PalmDB using kallisto translated search. Viral load obtained through alternative methods, such as RNA-ISH and qPCR, is compared to the target virus counts returned by kallisto. From left to right: 1. RNA-ISH (%) over total raw kallisto counts for SARS-CoV for 23 lung autopsy samples from COVID-19 patients obtained by bulk RNA sequencing^34^. Error bars show min-max values for each read in a pair; the dot shows the mean. 2. SARS-CoV-2 viral load by RT-qPCR (copies/mL) over total raw kallisto counts for SARS-CoV species obtained by bulk RNA sequencing of 16 saliva (circle), nasal swab (triangle), and throat swab (star) specimens from patients with acute SARS-CoV-2 infection^35,36^. Each specimen underwent duplicate library preparation and paired-end sequencing; points indicate the mean among the paired reads and duplicates, and error bars show min-max values. 3. Total raw kallisto counts for SARS-CoV species for 3 human iPSC-derived cardiomyocytes infected with SARS-CoV-2 and 3 control samples obtained by SMART-Seq^38^. 4. RT-qPCR (copies/mL) over total raw kallisto counts for ZEBOV for 19 rhesus macaque blood samples obtained during different stages of infection with ZEBOV and sequenced with Seq-Well^37^. **b**, To validate the mapping of nucleotide sequences to an amino acid reference with kallisto translated search and assess the robustness of the taxonomic assignment, we reverse translated all amino acid sequences in the PalmDB using the ‘standard’ genetic code (see Methods). The reverse translated PalmDB RdRP sequences were subsequently aligned to the optimized PalmDB amino acid reference (see Methods) with kallisto translated search. For each sequence, we differentiated the mapping result at each taxonomic rank into four categories: ‘correct’ or ‘incorrect’ taxonomic assignment based on the sOTU to virus ID mapping, ‘multimapped’ (the sequence aligned to multiple targets in the reference and could not unambiguously be assigned to one), or ‘not aligned’ (the sequence was not aligned). The plot shows the fraction of sequences falling into each mapping result category assessed at each taxonomic rank. The numbers above the bars indicate the total number of sequences per rank. Family names and numbers were omitted, and genera and species ranks were combined for readability.

**Fig. 4:**
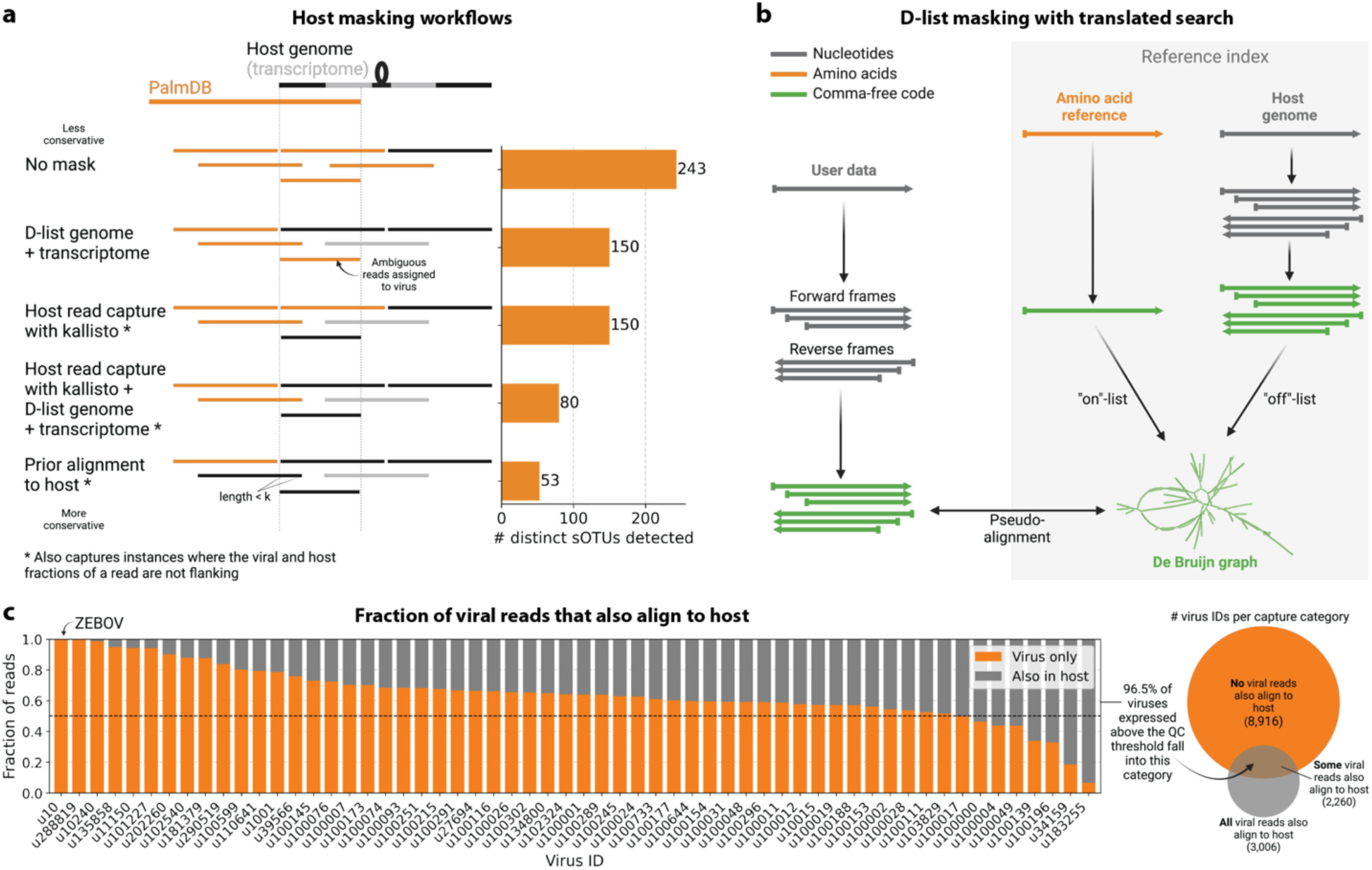
**a**, Schematic overview of the different host masking options discussed in this manuscript. Reads that align to PalmDB and are considered viral are marked in orange and reads that align to the host genome or transcriptome are marked in black or grey, respectively. The barplot shows the number of distinct sOTUs, defined by distinct virus IDs observed in ≥ 0.05 % of cells for each workflow. **b**, Schematic overview of masking the host genome with the D-list argument when used in combination with translated search. The D-list genome consists of nucleotide sequences and hence is translated to comma-free code in all six possible reading frames, similar to the translation of the nucleotide sequencing reads. **c**, Masking host sequences with the kallisto read capture workflow generates two distinct virus count matrices: The first contains viral reads that only aligned to the PalmDB, and the second contains viral reads that aligned to the host transcriptome in addition to the PalmDB. The majority of viruses detected above the quality control (QC) threshold (observed in ≥ 0.05 % of cells), had reads that aligned to the host transcriptome as well as the PalmDB. The barplot shows the fraction of reads for each virus that aligned to the PalmDB only (‘virus only’) and those that aligned to the host transcriptome in addition to the PalmDB (‘also in host’).

**Fig. 5:**
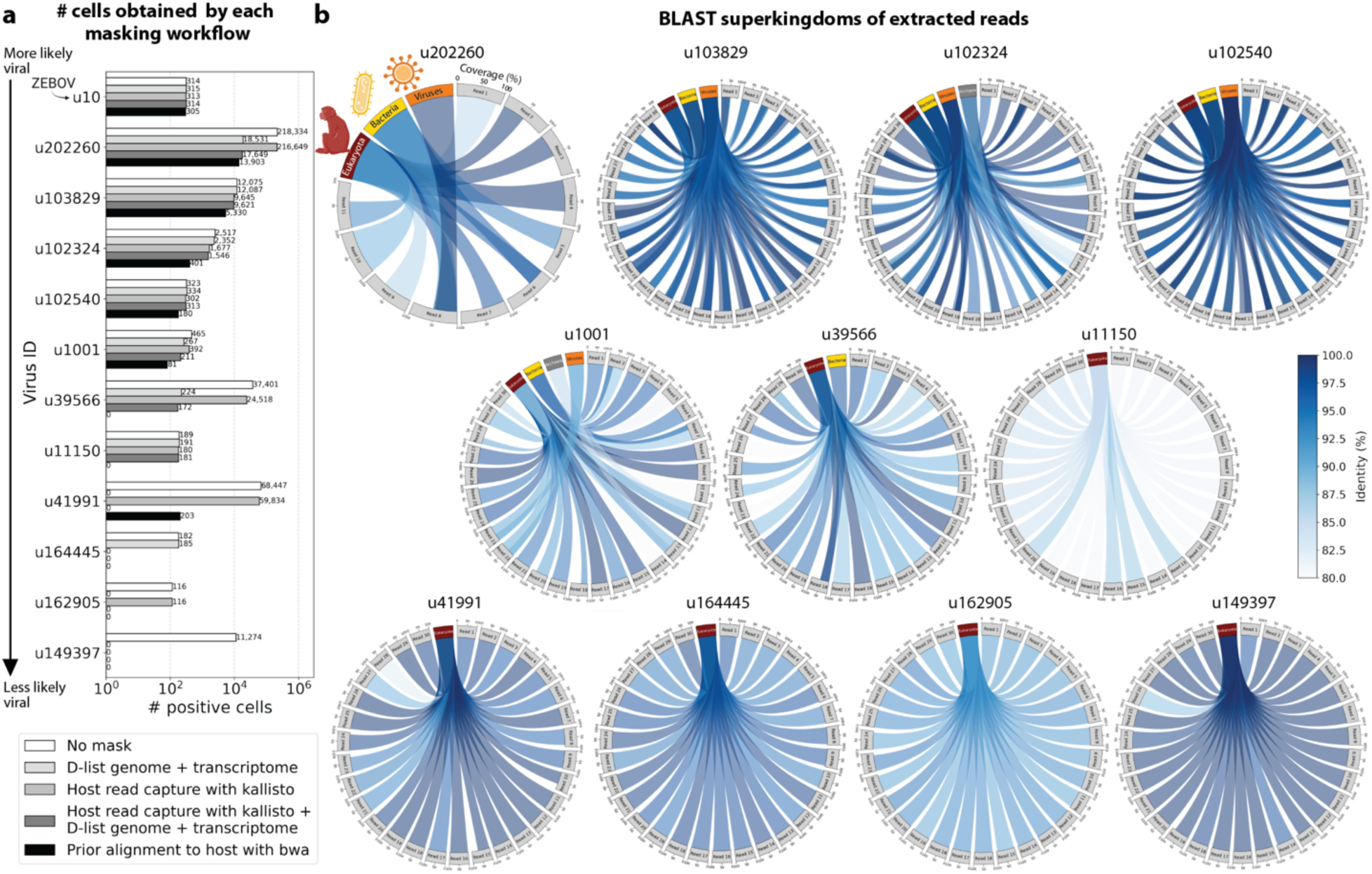
**a**, The number of positive cells obtained for 12 different virus IDs by each masking workflow. Each masking workflow is described in detail in the Methods section. The cell counts for all viruses detected above the QC threshold for all masking workflows are shown in Extended Data Fig. 5. **b**, pyCirclize plots showing the BLAST+^56^ results of randomly selected sequencing reads for each of the novel viruses shown in a (excluding the known virus ZEBOV which corresponds to virus ID ‘u10’). Each circular plot corresponds to the results for one virus ID. Each light grey sector corresponds to one sequencing read that links to the superkingdoms (red (eukaryotes), yellow (bacteria), orange (viruses), and dark grey (archaea) sectors) based on its BLAST+ alignment results. The width of the connecting link indicates the BLAST+ alignment coverage percentage, and its color indicates the identity percentage. For u202260, approximately ⅔ of the extracted reads yielded no BLAST results.

We first evaluated the impact of different host masking options on the resulting virome. We used kallisto translated search with PalmDB to map the virus profiles of peripheral blood mononuclear cell (PBMC) RNA sequencing samples from 19 rhesus macaques and applied different host masking workflows. The approach to masking host versus microbe reads and the handling of overlap between reference sequences can affect the downstream result. For example, sequences with varying sizes of virus-host overlap, sequences that span the junction of two exons, and entirely ambiguous sequences can influence the outcome of the masking and generate highly variable results depending on the method used (Extended Data Fig. 4a and 5). Depending on the research question and design, any one or a combination of different masking options might be appropriate. We explored the following masking options, listed from least to most conservative:

#### No mask

We aligned the sequencing reads to the PalmDB with kallisto translated search without masking or previously removing host sequences. For the macaque PBMC dataset, this masking option resulted in 243 distinct sOTUs detected (Fig. 4a).

#### D-list genome + transcriptome

To incorporate host read masking into our kallisto workflow, we quantified the reads while masking the host genome and transcriptome using an index created with the D-list (distinguishing list) option^53^. This option identifies sequences that are shared between a target transcriptome and a secondary genome and/or transcriptome. *k*-mers flanking the shared sequence on either end in the secondary genome are added to the index *de Bruijn* graph. During pseudoalignment, the flanking *k*-mers are used to identify reads that originated from the secondary genome but would otherwise be erroneously attributed to the target transcriptome due to the spurious alignment to the shared sequences. In our experiments, the target transcriptome consisted of the viral RdRP amino acid sequences contained in the PalmDB, and the secondary genome consisted of transcriptomic and genomic macaque and dog nucleotide sequences. When combining D-list with translated search, the secondary genome is translated to comma-free code in all six possible reading frames (Fig. 4b). This masking option can be easily added to the kallisto translated search workflow without any additional commands (Fig. 1). Masking the host transcriptome and genome with D-list resulted in 150 distinct sOTUs detected (Fig. 4a). Note that masking both the transcriptome and the genome, or either one will generate different results because masking only the genome will not mask sequences that span exon-exon junctions (Extended Data Fig. 4a and 5).

#### Host read capture with kallisto

To imitate prior alignment to the host genome, as performed with bwa (described below), within a simple, efficient kallisto workflow, we captured all reads that pseudoaligned to the host transcriptome with kallisto. Masking by capturing these host reads resulted in the same number of distinct sOTUs detected as masking with D-list (Fig. 4a).

#### Host read capture with kallisto + D-list genome + transcriptome

Although masking with D-list and capturing reads that aligned to the host transcriptome resulted in the same number of distinct sOTUs detected, the two methods masked different reads and resulted in different virus profiles (Fig. 5a, Extended Data Fig. 5). We decided to combine the D-list and host read capture masking approaches to achieve a conservative result similar to that achieved by prior alignment with bwa. In this approach, the sequencing reads were aligned to the PalmDB index with a D-list containing the host genome and transcriptome, and subsequently reads that pseudoaligned to the host transcriptome were captured. Combining the D-list and host read capture masking options reduced the number of detected sOTUs to 80 (Fig. 4a).

#### Prior alignment to host with bwa

We aligned the sequencing reads to the macaque and dog genomes using the highly sensitive alignment algorithm bwa^54^ and removed all reads that aligned anywhere in the host genomes before alignment to PalmDB with kallisto translated search. This achieved very conservative masking of the host genome. However, this workflow is complex, time-consuming, and computationally expensive (∼4.5 days using 60 cores for the macaque ZEBOV PBMC dataset). This workflow resulted in the detection of 53 distinct sOTUs (Fig. 4a).

There are inherent differences between these masking methods which are illustrated in Fig 4a and Extended Data Fig. 4a. Although the genome is passed to the software, the standard kallisto workflow builds an index based on the host transcriptome, not the entire host genome, since for genomes as large as the macaque genome, building the index on the entire genome would require a large amount of memory. Hence, masking by capturing reads that pseudoaligned to the host with kallisto will only capture host reads from mature mRNA molecules. If the D-list is passed both the transcriptome and the genome, it will be able to mask mature and nascent RNA molecules as well as RNA molecules originating from intergenic regions. The D-list index avoids excessive memory requirements by restraining the index to distinguishing sequences between viral and host sequences. As a result, reads that contain non-flanking host and viral sequences will not be filtered. Moreover, the D-list will favor viral assignment in the case of an entirely ambiguous read. Neither of these issues applies to masking with bwa since the alignment with bwa was performed against the host genome. Since bwa uses a smaller seed length than kallisto’s default *k*-mer size of 31, bwa provides more sensitive alignment of sequencing reads against the host genome and provides the most stringent filtering.

To confirm that reads identified as viral were not misaligned macaque reads, we extracted randomly selected sequencing reads from 11 virus IDs and aligned them against the nucleotide sequence database with BLAST+^56^ (Fig. 5b). The reads associated with virus IDs identified by all masking workflows (u202260, u103829, u102324, u102540, and u1001) BLAST-aligned with relatively low coverage and identity to several superkingdoms, including viruses. For u202260, approximately ⅔ of the extracted reads yielded no BLAST results. Given that the majority of RdRP sequences in the PalmDB originate from unknown viruses lacking reference genomes, it was expected that these sequences would not yield confident BLAST results. However, given the comprehensiveness of the macaque genome^57^, misaligned macaque sequences should BLAST to the macaque genome with high coverage and identity. The next two virus IDs, u39566 and u11150, were filtered out by the bwa workflow and did not BLAST to the viruses superkingdom. However, their BLAST results displayed relatively low coverage and identity, which would not be expected from macaque sequences. Below, we provide further evidence that u11150 sequences might have originated from an ongoing viral infection. This was likely an instance where filtering with bwa was too conservative and threw out viral sequences. u41991 was identified as viral by the bwa workflow but filtered out by the D-list + host capture workflow. Based on the BLAST results for u41991, which include high coverage and identity matches for eukaryotes, it is likely that filtering is the appropriate action. u164445 and u162905 were filtered by either capturing the host reads or using the D-list, respectively, and BLAST to eukaryotes with high coverage and identity, illustrating that a combination of the two methods leads to more robust results. Finally, sequences identified as u149397, which were filtered by all masking options and are only retained without masking, BLAST to eukaryotes with high coverage and identity.

Separately from exploring the results of different read masking options, we also investigated the question of virus filtering. Host read capture with kallisto generates two separate count matrices: One contains counts for reads that are solely viral, and a second contains counts for viral reads that also pseudoaligned to the host transcriptome. The distinction between filtering reads and filtering viruses becomes evident when examining the two count matrices: for the macaque PBMC dataset, we found that most viruses found in ≥ 0.05 % of cells had at least some reads that also mapped to the host transcriptome, including reads for ZEBOV (Fig. 4c and 5a). Moreover, aligning without host masking often led to the detection of more positive cells (Extended Data Fig. 5). Hence, naive masking of reads can lower the detection sensitivity of viruses that seem truly present. Our masking workflows facilitate the identification of viruses with a high likelihood of being truly present based on conservative host read masking, while also obtaining unmasked reads for these viruses to prevent the decrease in sensitivity inherent to masking host reads. We applied this approach when training the logistic regression models described below to minimize the occurrence of false viral absence.

#### The presence of novel putative viruses perturbs host gene expression in macaque blood cells, allowing prediction of viral presence based on host gene expression at single-cell resolution

We used kallisto translated search and the PalmDB to map the viral profiles of PBMC samples from 19 rhesus macaques sequenced at different stages of Ebola virus disease (EVD)^37^ (Fig. 6a) at single-cell resolution. The dataset consisted of 30,594,130,037 reads in total. After alignment to both the host genome (using the standard kallisto workflow) and PalmDB (using kallisto translated search with D-list + host capture masking), and quality control using the host count matrix (Extended Data Fig. 3a, Methods), we retained 202,525 PBMCs. We used the Leiden algorithm^58^ to partition the PBMC transcriptomes into 18 clusters of similar macaque gene expression, of which 16 could be assigned cell types based on common marker genes (Extended Data Fig. 3d).

**Fig. 6:**
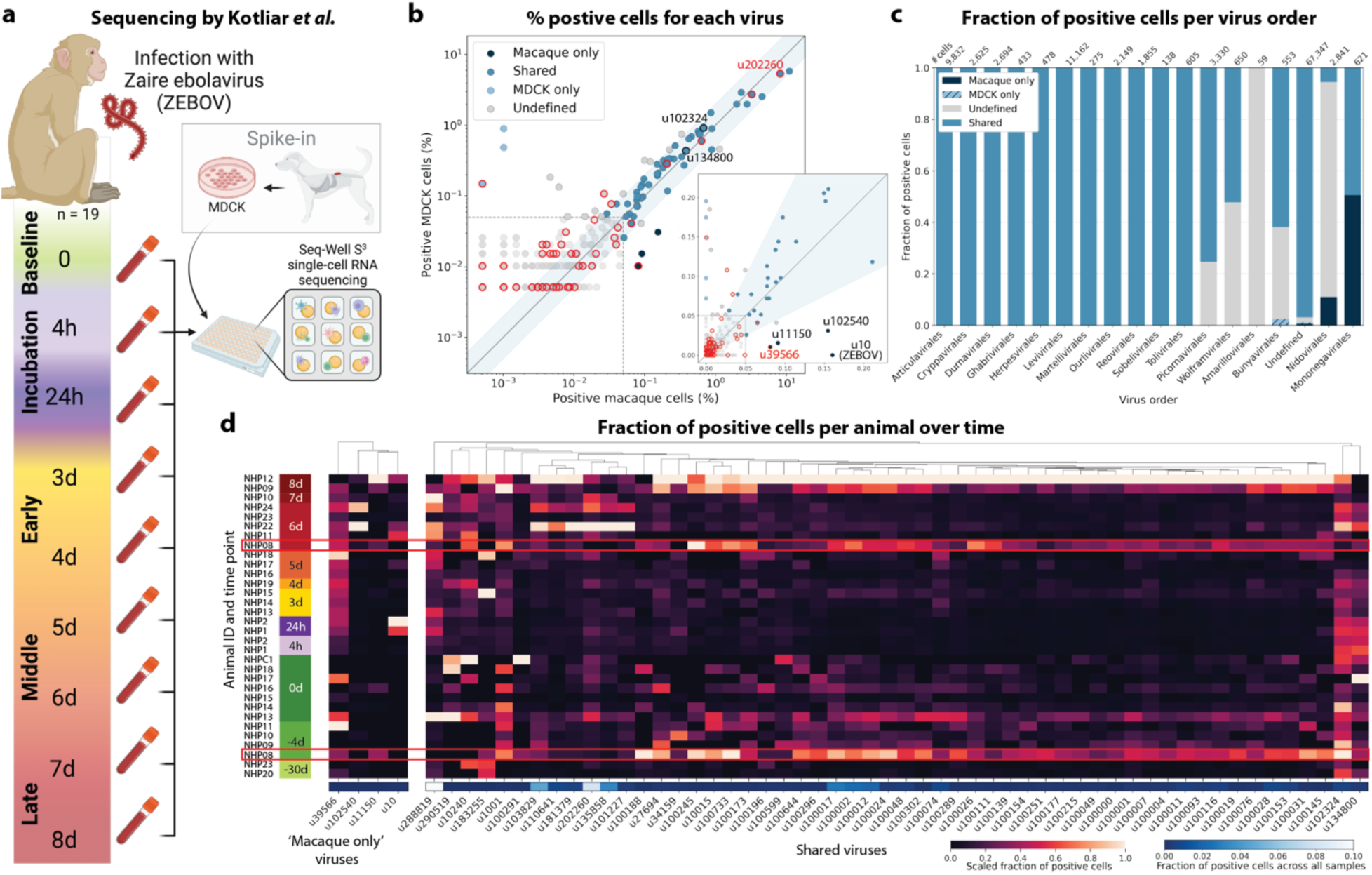
**a**, Schematic overview of the single-cell RNA sequencing data collected by Kotliar et al.^37^. Kotliar et al. performed single-cell RNA sequencing of peripheral blood mononuclear cell (PBMC) samples from 19 rhesus macaques at different time points during Ebola virus disease (EVD) after infection with Zaire Ebolavirus (ZEBOV) using Seq-Well^74^ with the S3 protocol^75^. A subset of the PBMC samples were spiked with Madin-Darby canine kidney (MDCK) cells. This schematic was adapted from the original design by Kotliar et al. **b**, For each virus-like sequence, the percentage of positive MDCK cells is plotted against the percentage of positive macaque cells. Virus IDs were categorized into ‘shared’, ‘macaque only’, ‘MDCK only’, and ‘undefined’ as described in the Methods section. The insert shows the same plot without log scale axes such that zero counts are included. A red edge marks contaminating virus-like sequences also observed in sequencing data obtained from blank sequencing libraries containing only sterile water and reagent mix (Extended Data Fig. 9c). **c**, Bar plot showing the fraction of positive cells obtained for each viral order, as defined by the PalmDB sOTUs, for each category. **d**, Fraction of positive cells for all ‘macaque only’ and ‘shared’ virus IDs. Each row corresponds to one animal at a specific EVD time point. The fractions were scaled to range from zero to one for each virus ID. The raw total fraction of positive cells for each virus ID across all samples is shown in blue below.

The obtained cell types, their marker gene expression and relative abundance over time are consistent with the results reported by Kotliar *et al.^37^*, including the emergence of a cluster of immature neutrophils and decreased lymphocyte abundance, especially natural killer cells, during EVD (Fig. 7a). While density based PBMC isolation typically removes neutrophils, immature neutrophils are less dense than mature neutrophils and can co-isolate with PBMCs during infections^37^. Clusters of the same cell type were often separated by time point (Fig. 7a), indicating changes in macaque gene expression within the same cell type over the course of the EVD. This is in agreement with results obtained by mass cytometry in Kotliar *et al.^37^*.

**Fig. 7:**
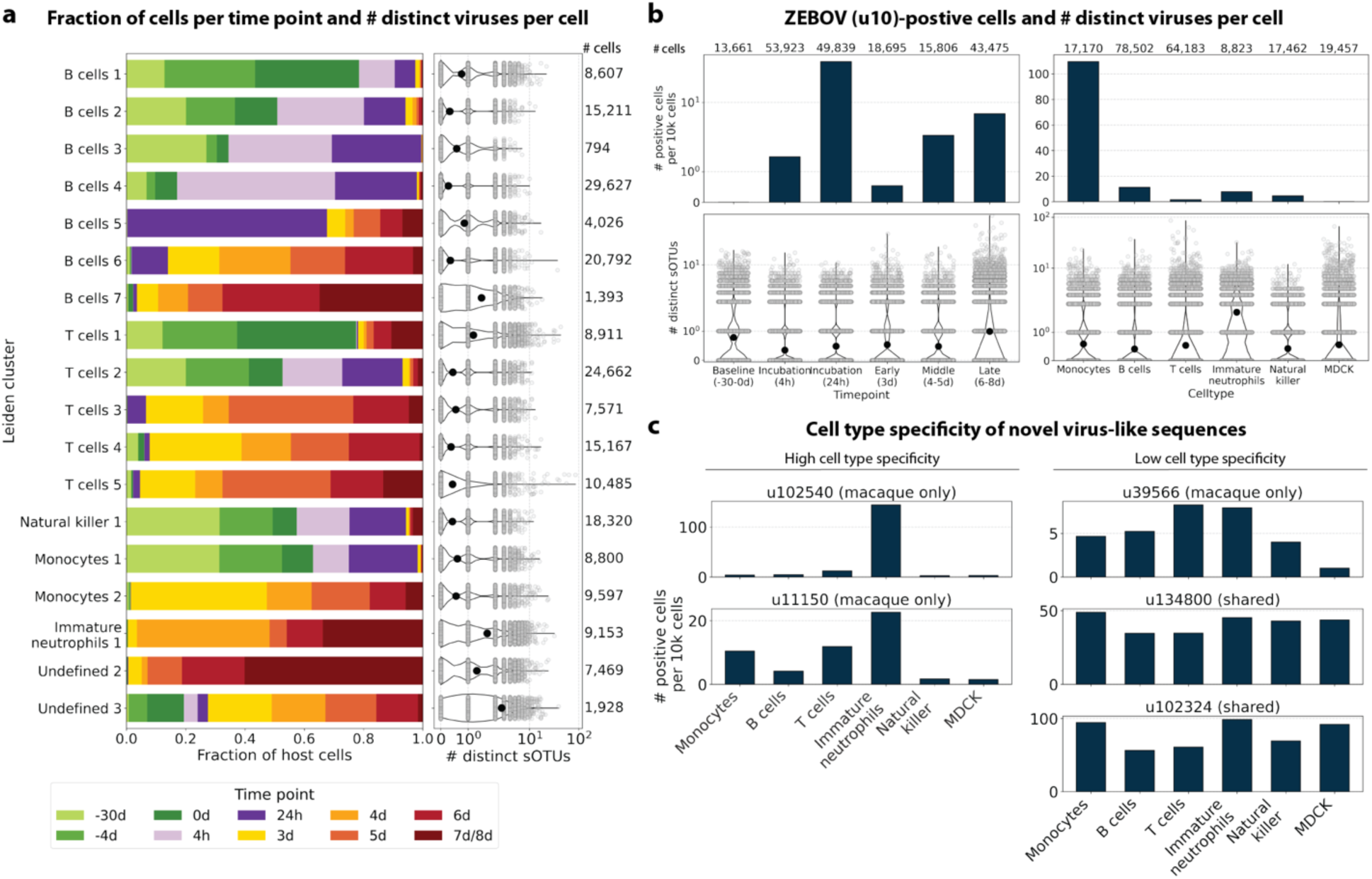
**a**, The fraction of cells occupied by each EVD time point is shown per Leiden cluster. Each Leiden cluster was assigned a cell type based on previously defined marker genes (Extended Data Fig. 3d). On the right, the number of distinct sOTUs detected in each cell is shown. Each grey dot corresponds to one cell, and the black dot corresponds to the mean across all cells. **b**, The number of ZEBOV (u10) positive cells per 10,000 cells is plotted per EVD time point (left) and per cell type (right). For each time point and cell type, the number of distinct sOTUs found per cell is plotted at the bottom. Each grey dot corresponds to one cell, and the black dot corresponds to the mean across all cells. **c**, The number of positive cells per 10,000 cells is shown per cell type for the remaining three (excluding ZEBOV) macaque only virus IDs and two shared virus IDs. Virus IDs that show relatively high cell type specificity are shown on the left, and virus IDs with relatively even detection across all cell types are shown on the right.

ZEBOV count data from this analysis workflow was also consistent with previously reported results. Since only a small fraction of the RNA molecules in these tissue samples are viral and of those, we only detect the RdRP, the measured absolute RNA counts for any one virus per cell are low (Extended Data Fig. 3e). As a result, we converted the virus count matrix into a binary matrix where each virus was recorded as being either present or not present in each cell. This approach has been previously validated for sparse single-cell RNA, specifically viral, sequencing data^24,59^, and prevented the need for further normalization by individual cellular viral load, which may introduce biases^49^. The presence of virus in each cell was then used to determine the viral abundance among populations of cells composed of clusters, cell types, or tissues. First, we used the binary virus count matrix to validate the detection of ZEBOV. Samples obtained during incubation displayed the highest abundance of ZEBOV-positive cells, and ZEBOV-positive cells remained detectable at all following time points (Fig. 7b, top left). These trends are consistent with the results reported by Kotliar *et al.^37^*.

The parallel analysis of viral and host gene counts at single-cell resolution allowed the identification of infected cell types based on host gene expression to reveal that ZEBOV-positive cells consisted predominantly of monocytes (Fig. 7b, top right). These results are consistent with previous literature on ZEBOV tropism^60^ and reproduce the ZEBOV abundance trends obtained by alignment to the ZEBOV genome^37^. This indicates that while the total viral counts obtained by kallisto translated search with PalmDB are low due to only detecting the RdRP, comparative trends are captured accurately. All *Ebolavirus* reads were identified correctly as ZEBOV with no counts detected for other *Ebolavirus* species (Extended Data Fig. 6).

Our analysis workflow identified virus-like sequences with sOTUs other than ZEBOV in this dataset. These virus-like sequences may be present due to, amongst others, viral infection of the host, host endogenous viral elements, infection of microbes residing in the host, infection of food ingested by the host, or laboratory contamination. Fig. 7b (bottom left and right) shows the total number of distinct sOTUs (corresponding to distinct virus IDs) detected over time and per cell type. We observed a slight increase in the number of distinct sOTUs detected per cell during the later stages of EVD, driven by T cell, B cell, and neutrophil clusters with high fractions of cells during later EVD stages (Fig. 7a). Neutrophils showed the highest numbers of distinct sOTUs per cell (Fig. 7b, bottom right). Since neutrophils fulfill their microbicidal function through phagocytosis and pinocytosis, it is possible that viral RNA was picked up by these cells through ingestion. In the following paragraphs, we explore different approaches to interpret the presence of these virus-like sequences.

Among the samples in this dataset, we detected a total of 11,176 virus-like sequences with at least one read that aligned to the PalmDB and did not align to the host (Fig. 4c), including many sOTUs from genera known to infect rhesus macaques (Extended Data Fig. 6)^61^. However, the majority of these virus-like sequences were expressed in less than 0.05 % of cells, which we defined as a quality control (QC) threshold (Fig. 6b, Methods). All of the virus-like sequences with positive cell fractions above the QC threshold in macaque cells that passed quality control can be explored in this interactive Krona plot^62^ broken down by animal, timepoint, taxonomy, and fraction of cells occupied by each virus: https://tinyurl.com/23h6k36u.

A subset of samples included a spike-in of Madin-Darby canine kidney (MDCK) cells, resulting in a total of 23,500 MDCK cells after quality control and species separation (Extended Data Fig. 3b and c, Methods). We used the spike-in to further break down the viruses into ‘macaque only’, ‘MDCK only’, and ‘shared’ viruses (Fig. 6b, Methods). We expected that shared viruses occurring in both macaque and MDCK cells would include viruses introduced by the contamination of laboratory reagents used during sample preparation and sequencing^63^, cell-free RNA contamination, endogenous retroviruses^52,64^, and widespread latent infections. After filtering and categorization of viruses, we detected 4 (including ZEBOV) macaque only, 7 MDCK only, 15 undefined, and 54 shared viruses. This result suggests that the majority of virus-like sequences detected in this dataset were introduced through contamination. Indeed, many virus-like sequences that fell into the shared category could also be detected in ‘blank’ sequencing libraries containing only sterile water and reagent mix, providing evidence for their origin in widespread laboratory reagent contamination^63^ (Fig. 6b, Extended Data Fig. 9c). The sOTUs of the macaque only and shared virus IDs, when available, are listed in Extended Data Table 1. Fig. 6c shows the fraction of reads occupied by each viral order (as defined in the sOTU for each virus ID) for macaque only, MDCK only, and shared viruses. Among viruses shared between macaque and MDCK cells, *Levivirales* (recently renamed to Norzivirales^65^), *Articulavirales* (which include the family of influenza viruses), and viruses of unknown taxonomy made up the largest fractions. *Norzivirales* are an order of bacteriophages, the majority of which were discovered in metagenomics studies^66^. They might have been introduced through bacterial contamination during sample preparation and sequencing. The shared viruses also included orders such as *Herpesvirales*, which are widespread, sometimes spreading through cross-species infections, and are known to persist in their host as latent infection^67,68^. Virus-like sequences detected in MDCK cells included sOTUs from the order of *Bunyavirales*, which infect a wide range of hosts, including MDCK cells^69^, as well as virus-like sequences of unknown order. Virus-like sequences found only in macaque cells were of unknown order, in the order *Mononegavirales*, which includes ZEBOV, and in the order *Nidovirales*, which are known to infect mammals and include the family *Coronaviridae*. Virus-like sequences of known order (based on the sOTU) for each group are reasonably expected to be present in the respective sample types and the context of the hosts, which supports the biological validity of these viral read classifications.

To visualize the virus profiles of individual animals and over time, we plotted the fraction of positive cells for each macaque only and shared virus ID per animal and time point (Fig. 6d). The relative viral abundances varied, both between individual monkeys and time points. Notably, in some instances where the same animal was measured across several time points, the viral profile of this animal was reproduced in the later time point (Fig. 6d, Fig. 8a). The viral profiles of animal NHP08 at -4 days pre-infection and 6 days post-infection with ZEBOV are highlighted in the heatmap (Fig. 6d). Animals NHP1 and NHP2 each had two samples sequenced 20 hours apart which displayed highly similar viral profiles for each animal over time (Fig. 8a). This suggests that viral profiles sampled and sequenced within a short time window are coherent over time and across samples which is consistent with expectation. Several virus-like sequences, including u102324, were present in all animals and time points with relatively similar abundance (Fig. 6d, Extended Data Fig. 7a), coherent with their classification as shared sequences likely originating from contamination.

**Fig. 8:**
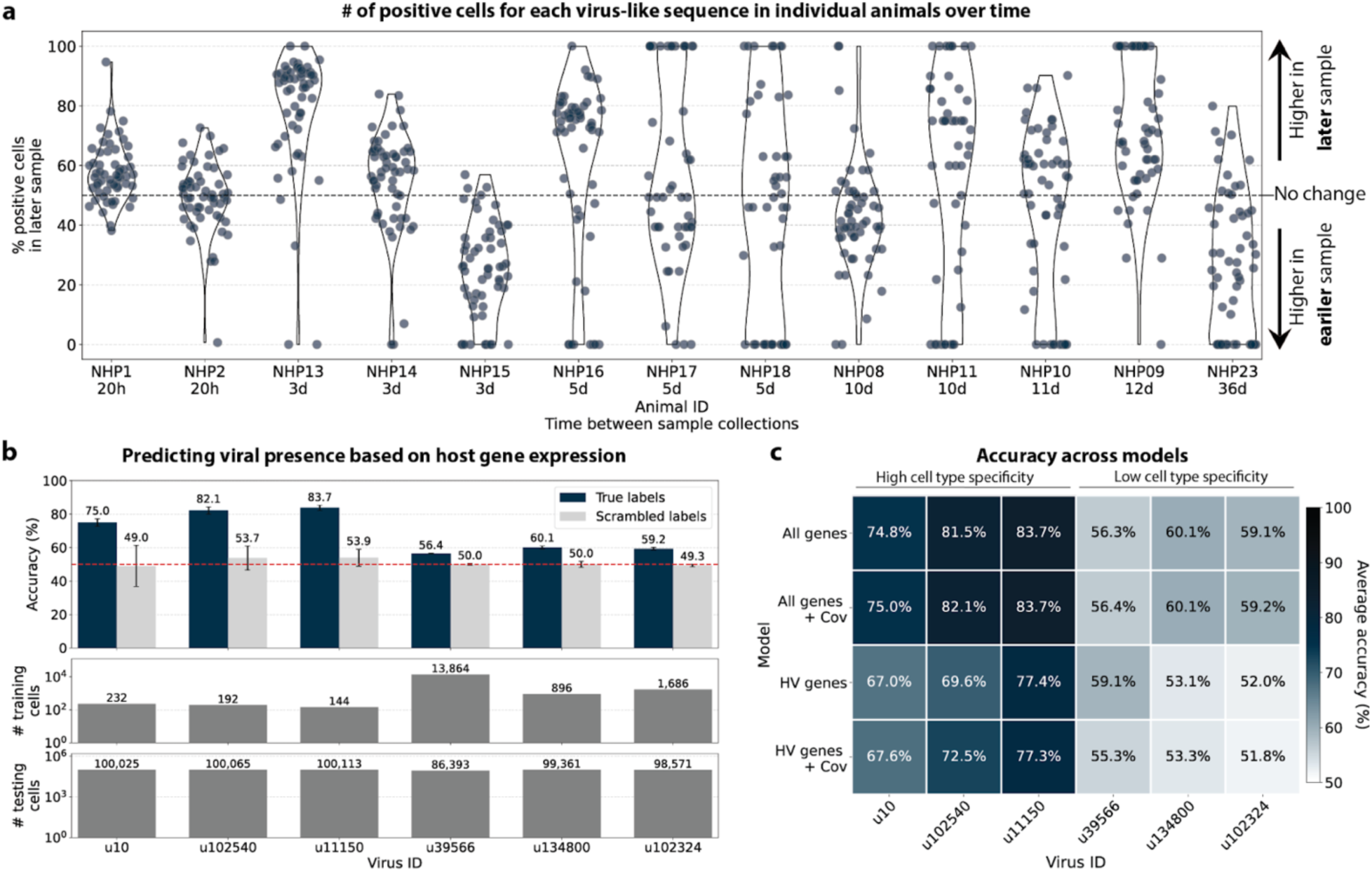
**a**, Several animals included in the macaque PBMC dataset were sampled twice, at two different time points. Here, for each virus-like sequence, the percentage of positive cells occupied by the later time point is shown. The number of positive cells for each virus-like sequence was first normalized to the total number of cells in the sample. Only virus-like sequences for which at least one time point had positive counts were included for each animal. A percentage of 50% indicates that the number of positive cells for that virus-like sequence remained stable between the two time points. **b**, We trained logistic regression models to predict the presence of specific virus-like sequences based on host gene expression at single-cell resolution. The accuracy of the logistic regression model trained on all macaque genes with donor animal and EVD time point as covariates is shown for the known virus ZEBOV (u10) and five novel virus IDs. The presence of virus-like sequences that displayed high cell type specificity could be predicted with >70 % accuracy, while virus-like sequences with low cell type specificity could not be predicted above random chance (50 %, marked by the red dashed line). As a negative control, viral presence and absence labels were scrambled at random in the training data, causing the prediction accuracy to drop to random chance (50 %), as expected. The error bars indicate the standard deviation between models initialized with different random seeds. The bottom bar plots show the number of testing and training cells for each virus (also see Extended Data Fig. 8c). **c**, Heatmap of the prediction accuracy (averaged across models initialized with different random seeds) across all possible modeling combinations (training on all macaque genes versus only highly variable (HV) genes, and with or without covariates donor animal and EVD time point).

We then attempted to further determine which virus-like sequences were likely present due to viral infection of the host based on cell-type specificity and a coinciding host antiviral response. We visualized the viral tropism of the remaining three (other than ZEBOV) macaque only viruses. Two of them, u102540 and u11150, displayed relatively high sample-specificity while u39566 was abundant across all samples, similar to the shared viruses u134800 and u102324 (Extended Data Fig. 7a). The sOTU of u102540 indicates that it is an *Alphacoronavirus sp.*, which are known to infect rhesus macaques^61^. u102324 is predicted to belong to the family *Iflaviridae* (Extended Data Table 1), which is a family of viruses that infect insects^70^, and the viral reads from this virus ID were likely not the result of an ongoing viral infection. The remaining virus IDs, u11150, u39566, and u134800, are of unknown taxonomy across all taxonomic ranks.

Two virus-like sequences exhibited cell-type specificity suggestive of viral infection. Of the three macaque only virus IDs excluding ZEBOV, we found that u102540 and u11150 showed high cell type specificity, while u39566 was expressed more evenly across all cell types (Fig. 7c). While u39566 was categorized as ‘macaque only’ above, it is likely a contaminating sequence given its presence in the blank sequencing libraries (Extended Data Fig. 9c). The lack of cell-type specificity coincides with u39566 sequences originating from reagent contamination and illustrates the importance of combining several different approaches, as described here, when interpreting the presence of virus-like sequences. u102540 (*Alphacoronavirus sp.*) exhibited high fractions of positive cells in neutrophils, while u11150 also displayed lower expression in monocytes, B cells, and T cells. Neutrophils play an important role in the innate immune response and promote virus clearance through phagocytosis. During phagocytosis, neutrophils engulf virions and apoptotic bodies. It is possible that the cell type specificity towards neutrophils observed here was due to neutrophils engulfing viral RNA during phagocytosis rather than viral tropism. As expected, the shared viruses u134800 and u102324 did not display cell type specificity (Fig. 7c).

The simultaneous analysis of the host and virus count matrices supported that several viruses identified were likely infecting the host and revealed virus-induced host gene expression. We hypothesized that viral presence in individual cells may be predicted based on the host gene expression. Since our workflow maintains single-cell resolution, we can analyze viral presence and host gene expression at single-cell resolution in parallel and investigate whether the presence of a virus affects host gene expression. We trained logistic regression models for all macaque only and shared (present in both MDCK and macaque cells) virus-like sequences to predict viral presence or absence in individual cells based on the cell’s host gene expression. The models were either trained on all or only highly variable macaque genes and with or without the donor animal and time point as covariates. After training models using a random selection of virus-positive and an equal number of virus-negative cells, we tested the model predictions on held-out test cells (Fig. 8b, Extended Data Fig. 8a and e). Given the cell type specificity of several of the virus-like sequences, virus-negative training cells were selected to be of the same cell types as virus-positive cells to ensure that we were not simply predicting cell type rather than viral presence.

We found that the presence or absence of virus-like sequences that displayed cell type- and sample-specificity (u10 (ZEBOV), u102540, and u11150) could be predicted at > 70 % accuracy (Fig. 8b and c), although for u11150, the sensitivity decreased with the inclusion of the covariates donor animal and EVD time point (Extended Data Fig. 8a). The sensitivity and specificity are shown in Extended Data Fig. 8a. By contrast, the presence of viruses that did not display cell type-specificity (u39566, u134800, and u102324) could not be predicted better than random chance (50 %) (Fig. 8b and c, Extended Data Fig. 8a). As a negative control, we scrambled the binary virus count matrix used for model training, effectively randomizing the presence or absence of a virus in each cell. As expected, the prediction accuracies dropped to those expected at random (50 %) (Fig. 8b and c). We also confirmed that the different virus-like sequences with high prediction accuracy, including the known infection with ZEBOV (u10), were not present in the same cells (Extended Data Fig. 7b).

This learnable relationship between viral presence and host gene expression provides further evidence that reads from u10, the known infection with ZEBOV, as well as novel virus-like sequences u102540 and potentially u11150, originated from an ongoing viral infection and/or viral clearance which perturbed host gene expression at the single-cell level.

The only other virus-like sequence which displayed prediction accuracies > 70% was u202260 (Extended Data Fig. 8e). This was surprising, as u202260 was categorized as shared between macaque and MDCK cells and was also present in the blank sequencing libraries (Fig. 6b, Extended Data Fig. 9c), indicating that it likely originated from laboratory reagent contamination. However, although its prediction accuracies were relatively high, the gene weight correlation between different models was low for u202260 (Extended Data Fig. 8b) and the standard deviation of gene weights within the same model generated with different random seeds was comparatively high (Extended Data Fig. 9a), indicating that genes were weighted differently between models and seeds for u202260. This suggests that a shared feature across genes, such as cell health or sequencing depth, was learned rather than the expression of specific genes.

To explore virus-induced host gene expression, we identified macaque genes with the largest predictive power and smallest variation (across models initialized with different random seeds), for the regression models trained on highly variable genes with the donor animal and time point as covariates (Extended Data Fig. 9a). Approximately one third of the macaque Ensembl IDs did not have annotated gene names, which is a common problem for genomes from non-model organisms. We used gget^45^ to translate annotated Ensembl IDs to gene symbols and to perform an enrichment analysis on the returned gene symbols using Enrichr^71^ against the 2023 Gene Ontology (GO) Biological Processes database^72^. The highly weighted genes for u10 (ZEBOV) returned significant enrichment results for several virus-associated GO terms including ‘Negative Regulation Of Viral Entry Into Host Cell (GO:0046597)’, ‘Negative Regulation Of Viral Life Cycle (GO:1903901)’, and ‘Regulation Of Viral Entry Into Host Cell (GO:0046596)’, validating our approach for the identification of genes associated with a virus-related host gene response. Similarly, the enrichment analysis of highly weighted genes for the novel virus ID u11150 mapped to ‘Receptor-Mediated Endocytosis Of Virus By Host Cell (GO:0019065).’ For virus ID u102540, several highly ranked GO terms were indicative of an inflammatory response, such as ‘Positive Regulation Of Type II Interferon Production (GO:0032729)’ and ‘Positive Regulation Of Cytokine Production (GO:0001819)’. Several predictive genes were associated with the positive regulation of cytokine production and modulation of inflammation (e.g., FCN1 for u 10, MAPK11 for u11150, and CD14 for u102540). Overall, these results provide further evidence that the novel virus-like sequences u102540 and u11150 originated from an ongoing viral infection or clearance resulting in a host gene response.

## Discussion

Our work provides a method for extracting a ‘virome’ modality from any bulk or single-cell RNA-seq data by leveraging a new method that maps and quantifies species-level viral RdRP sequences against an amino acid reference. We built on the existing alignment software kallisto^44^ and bustools^76^ and expanded them for translated alignment by (reverse) translating both the amino acid reference and the nucleotide sequencing reads into a common, nonredundant comma-free code. While we validated kallisto translated search in combination with PalmDB for the identification of viral RNA, our novel workflow can be applied in combination with any amino acid reference. kallisto translated search permits the alignment of nucleotide sequencing data to any amino acid reference at single-cell resolution. For example, amino acid sequences of antimicrobial peptides^77^ can be used as a reference to identify these peptides in bulk and single-cell RNA sequencing data. Moreover, amino acid transcriptomes of homologous species may be used as a reference for species with missing or incomplete reference genomes. In this case, operating in the amino acid space will increase similarity due to the robustness to single-nucleotide mutations.

We validated kallisto translated search in combination with PalmDB for the detection and identification of viral RNA in next-generation sequencing data at single-cell resolution. As we noted in the introduction, the number of viruses expected to cause human infectious disease is eclipsed by the comparatively few viruses with complete reference genomes and the even smaller number of viruses that have been detected in humans. It is important to monitor the presence of viruses in the human population, both to prevent pandemic outbreaks and to further understand the role of viruses in various diseases. We have shown that such monitoring and novel virus discovery can be performed using single-cell RNA-seq data. Moreover, our work provides a platform for characterizing omnipresent virus-like sequences associated with different environments, hosts, and laboratory reagents.

The virus count matrix, which is obtained using kallisto translated search in combination with PalmDB, is an entirely new modality that we have begun to explore in this paper. We found that this matrix is sparse with relatively low molecule counts per cell (Extended Data Fig. 3e). While using the highly conserved RdRP to identify viruses makes our workflow very efficient and is the key to being able to detect over 100,000 distinct viruses, RdRP RNA only makes up ∼1 % of the total viral RNA present in the sequencing data analyzed here (Extended Data Fig. 2) resulting in the sparsity of the virus count matrix. We anticipate that this number varies between virus species and sequencing technology, making it difficult to define a general detection limit. To normalize this sparse and low-count matrix, we binarized the virus count matrix such that each cell was either positive or negative for each virus. Given the low counts, we expect that there is a high occurrence of false negatives in the virus count matrix while the confidence in positive cells is high. However, we have shown that relationships between viral presence and host gene signatures can be learned regardless.

A common problem in the identification of microbial sequences is the misidentification of host sequences as microbial. The PalmDB is not a curated database, and it is possible that some virus-like sequences in the PalmDB are not derived from viruses. In addition, differentiating between ongoing infections, reagent or sample contamination, cell-free RNA contamination, endogenous retroviruses, and widespread latent infections is a challenge. The kallisto translated search method computes both the virus count matrix and the host gene expression matrix at single-cell resolution, providing unique opportunities for parallel analysis of viral signatures and their effect on host gene expression. We describe different approaches to evaluate the nature of viral sequences identified by kallisto translated search, including taxonomic assignment of viruses based on the sOTU, analysis of viral tropism, extraction and BLAST alignment of raw sequencing reads identified as viral, and using a sample spike-in to categorize viruses into shared and sample-specific viruses. Moreover, we describe and evaluate different workflows to mask the host genome and/or transcriptome, allowing different levels of conservativeness and the quantification of sequencing reads that align to viral RdRPs as well as the host transcriptome. Notably, the efficacy of masking the host genome and/or transcriptome will depend on the quality and comprehensiveness of said genome/transcriptome. In this case, the majority of host sequences originated from rhesus macaque, which has a very comprehensive genome assembly^57^. Finally, we trained logistic regression models to predict viral presence at the single-cell level based on host gene expression, achieving high accuracy indicative of an ongoing viral infection or clearance. Our results show that it is beneficial to combine multiple of these approaches, which we validate and describe in detail, for the interpretation of the presence of virus-like sequences.

Focusing on the RdRP produces biases between virus species with varying life cycles, depending on the sequencing technology used. The genome of many negative-strand RNA (-ssRNA) is replicated as well as transcribed. Transcription produces short, often polyadenylated mRNA products which are captured and sequenced, including the RdRP. In contrast, the genome of many positive-strand RNA (+ssRNA) viruses undergoes replication, but not transcription. Instead, the genome is translated into polyproteins, which are subsequently cleaved. While +ssRNA virus genomes are often polyadenylated and hence are captured by polyA capture-dependent single-cell RNA sequencing technologies, sequencing ∼100 bases from the polyA-tail using a poly(T) primer will not capture the RdRP if it is located too far from the polyA-tail (see the schematic overview of the SARS-CoV genome in Fig. 1). In this scenario, the RdRP of +ssRNA viruses will, however, be captured by bulk RNA sequencing and random hexamer primers in single-cell RNA sequencing (Extended Data Fig. 4c). Hence, sequencing using random hexamer primers overcomes the virus life cycle-dependent bias for single-cell technologies. Many novel sequencing technologies, including Parse Biosciences SPLiT-Seq^33^, employ random hexamer primers to produce full-coverage sequencing and overcome biases introduced by poly(T) primers. We foresee that the use of random priming in sequencing will continue to increase. It is worth noting that, depending on the technology, intra-genomic sequences of +ssRNA viruses might be captured by poly(T) primers nonetheless due to mispriming. Even with random priming, many biases will remain. For example, any viral RNA that is not polyadenylated will not be captured efficiently by single-cell sequencing technologies that rely on polyA capture.

We hope that kallisto translated search will be widely implemented in the analysis of next-generation sequencing data to identify the presence of viral RNA, as well as inform the experimental design of research aiming to identify microbial reads from RNA sequencing data. We describe several experimental design choices that greatly impact the results of microbial read quantification, such as the sequencing primer design and sample spike-ins. The masking workflows described in this paper and the associated challenges are applicable to any metagenomics analysis beyond the identification of viral reads, and the workflows described here can be easily applied to nucleotide references, such as a 16S database for the characterization of the human gut microbiome^78^.

## Methods

### Developing kallisto translated search and optimization for the identification of viral RNA

#### Building kallisto translated search and choosing a new ‘genetic code’

To perform translated alignment, the nucleotide and amino acid sequences need to be translated into a shared ‘language’. This might be achieved by translating nucleotides to amino acids or vice versa. Since kallisto encodes nucleotide characters in 2 bits (allowing a total of 4 distinct nucleotides to be encoded), encoding the 20 different amino acids resulting from translated nucleotide sequences was not feasible. Moreover, reverse translating the amino acid sequences to nucleotides would be intractable due to the redundancy in the genetic code leading to a combinatorial explosion in nucleotide sequences consistent with an amino acid reference. We therefore translated the nucleotide sequences and reverse translated the amino acid sequences using a fixed synthetic code designed to reduce spurious alignments. We explored two different codes for this translation: 1. Comma-free code and 2. A code that maximizes the Hamming distance between frequently occurring amino acids (Extended Data Fig. 10a). While maximizing Hamming distance is advantageous in terms of avoiding sequence multimapping (see next paragraph), a comma-free code prevents off-frame alignment since any k-mers formed by adjacent words will not be included in the dictionary. We found that the comma-free code recalls viral sequences equally well compared to maximizing the Hamming distance between amino acids (Extended Data Fig. 10b).

#### Optimization of PalmDB for the identification of viral reads in RNA sequencing data

Due to the occurrence of the ambiguous amino acid characters B, J, and Z, 62 out of 296,623 viral sequences were transformed into identical sequences after reverse translation to comma-free code. The identical sequences were merged and assigned a representative virus ID. Due to the high similarity between viral RdRP sequences, the loss of aligned sequences due to multimapping to several reference sequences was a major concern. Moreover, the necessity of reverse translating the amino acid sequences further decreases the Hamming distance between reference sequences by approximately 30 % (Extended Data Fig. 10d). To overcome this problem, we tried clustering the amino acid sequences based on 99 % similarity using the MMseqs2 algorithm^79^. This resulted in 6,518 clusters with high concordance of taxonomy labels between sequences in the same cluster (Extended Data Fig. 10e). However, although clusters were computed correctly based on their concordance with taxonomy, this resulted in 67.4 % of sOTUs not being detected anymore (compared to 3.3 % when using the complete index). As a result, we decided to group the sOTUs instead, treating virus IDs with the same taxonomy across all main taxonomic ranks like transcripts of the same gene (available here: https://tinyurl.com/4wd33rey). This retained the alignment percentage of the complete index while allowing highly accurate taxonomic assignment and minimal sequence loss to multimapping (Fig. 3b). It is noteworthy that the default kallisto k-mer length of 31 nucleotides equals only 10 amino acids. Given the architecture of the current kallisto version (0.50.1), which is optimized for 64-bit k-mers with each nucleotide occupying two bits, k cannot be set > 31. This will change in future versions.

### Validation and benchmarks

#### Visualization of the Kraken2 and kallisto translated search alignments of ZEBOV sequences

Kotliar *et al.^37^* performed single-cell RNA sequencing of PBMC samples from rhesus macaques after infection with ZEBOV (further described below). A subset of the data obtained by Kotliar *et al.* at 8 days post-infection with ZEBOV was used to visualize the identification of RdRP sequences using kallisto (v0.50.1) translated search. The first 100,000,000 raw sequencing reads from the <u>GSE158390</u> library SRR12698539 were aligned to the ZEBOV reference genome (NC_002549.1) using Kraken2 v2.1.2 and to the optimized PalmDB using kallisto translated search. Aligned reads from both workflows were extracted and realigned to the ZEBOV genome using bowtie2^39^ v2.2.5 and SAMtools^40^ v1.6 as previously described^80^. The visualization shown in Extended Data Fig. 2 was generated from the resulting sorted bam files with the NCBI Genome Workbench^41^.

#### Testing robustness to mutation

676 *Zaire ebolavirus* (ZEBOV) RdRP sequences were identified by aligning the first 100,000,000 raw sequencing reads from the <u>GSE158390</u> library SRR12698539 to the optimized PalmDB using kallisto translated search. Mutation-Simulator^48^ (v3.0.1) was used to add random single nucleotide base substitutions to the RdRP sequences at increasing mutation rates. We performed 10 rounds of simulated mutations per mutation rate. The sequences were subsequently aligned using kallisto translated search against the complete PalmDB, Kraken2 translated search against the RdRP amino acid sequence of ZEBOV with a manually adjusted NCBI Taxonomy ID to allow compatibility with Kraken2, and kallisto standard workflow against the complete ZEBOV nucleotide genome (GCA_000848505.1). We subsequently calculated the recall percentage over all 676 sequences. For kallisto translated search, the recall percentage was calculated based on species-level taxonomic assignment. Since the other two methods were only given the target virus sequence as a reference and did not have to distinguish between different viruses, their recall percentage was calculated based on all aligned sequences. The recall percentage over all 676 sequences for the 10 rounds at each mutation rate is shown in Fig. 2c. Extended Data Fig. 4b shows the precision with which kallisto translated search identified the correct virus versus other taxonomies at each mutation rate. The recall and precision at mutation rates > 0 were fitted with an inverse sigmoid function using non-linear least squares using the scipy.optimize.curve_fit function (scipy v1.11.1).

#### Alignment and quantification of viral counts in validation datasets

The sequencing reads for each library used in the validation (Fig. 3a) were aligned with kallisto translated search against the PalmDB index D-listed with the corresponding host genome and transcriptome. The hosts were (i) human (GRCh38 Ensembl version 109) for <u>GSE150316</u>, <u>GSM4548303</u> and the SARS-CoV-2 saliva, nasal, and throat samples, (ii) mouse (GRCm39 Ensembl version 109) for <u>GSM5974202</u>, and (iii) rhesus macaque (Mmul_10 Ensembl version 109) and dog (ROS_Cfam_1.0 Ensembl version 109) for <u>GSE158390</u>. Count matrices were generated with bustools (v0.43.1). Fig. 3a shows the total raw counts obtained for each target virus species. RT-qPCR and RNA-ISH counts were reproduced from the original publications.

#### Validating the alignment of nucleotide sequences to an amino acid reference and assessing the accuracy of the taxonomic assignment

To validate the mapping of nucleotide sequences to an amino acid reference with kallisto translated search and assess the accuracy of the taxonomic assignment, we reverse translated all amino acid sequences in the PalmDB using the ‘standard’ genetic code from the biopython^81^ (v1.79) Bio.Data.CodonTable module and DnaChisel^82^ (v3.2.10) (with a slight adaptation to allow the ambiguous amino acids ‘X’, ‘B’, ‘J’, and ‘Z’ occurring in the PalmDB, which was later implemented in DnaChisel v3.2.11). A unique synthetic ‘cell barcode’ was generated for each resulting nucleotide sequence, and the sequences were aligned to the optimized amino acid PalmDB with kallisto translated search, keeping track of each sequence separately as if they were an individual cell. The synthetic barcodes allowed subsequent analysis of the alignment result for each individual sequence, and the accuracy of the obtained taxonomy based on the virus ID to sOTU mapping provided by PalmDB is shown in Fig. 3b. For each sequence, we differentiated between ‘correct’ or ‘incorrect’ taxonomic assignment, or, if the sequence did not return any results, whether it was ‘multimapped’ (the sequence aligned to multiple targets in the reference and could not unambiguously be assigned to one) or ‘not aligned’ (the sequence was not aligned), at each taxonomic rank.

### Analysis of macaque PBMC data

Kotliar *et al.^37^* performed single-cell RNA sequencing of PBMC samples from 19 rhesus macaques at different time points during Ebola virus disease (EVD) after infection with ZEBOV (EBOV/Kikwit; GenBank accession MG572235.1; *Filoviridae: Zaire ebolavirus*) using Seq-Well^74^ with the S3 protocol^75^. A subset of PBMC samples were spiked with Madin-Darby canine kidney (MDCK) cells. The data is available at <u>GSE158390</u>, and we obtained the raw sequencing data from the European Nucleotide Archive using FTP download links and ffq (v0.3.0)^83^. The data is split into 106 datasets containing 30,594,130,037 reads in total.

#### Alignment to the host transcriptome

The rhesus macaque Mmul_10 and domestic dog ROS_Cfam_1.0 genomes were retrieved from Ensembl version 109. The reference index was built using both genomes and the kb-python (v0.28.0 with kallisto v0.50.1 and bustools v0.43.1) ref command to create a combined index containing the transcriptome of both species. We quantified the gene expression in each of the 106 datasets using the standard kallisto-bustools workflow^13^ with the ‘batch’ and ‘batch-barcodes’ arguments to process all files simultaneously while keeping track of each batch, and with the ‘x’-string ‘0,0,12:0,12,20:1,0,0’ to match the Seq-Well technology. Since the Seq-Well technology does not provide a barcode on-list, we generated a barcode on-list using the ‘bustools allowlist’ command, requiring each barcode to occur at least 1,000 times. We subsequently corrected the cell barcodes using the generated on-list and computed the count matrix using the ‘bustools count’ function.

#### Host cell quality control, filtering, and separation of macaque and MDCK cells

The count matrix generated by bustools was converted to h5ad using kb_python.utils.kb_utils and read into Python using anndata v0.8.0. Metadata such as donor animal, the presence of an MDCK spike-in, and time point were added to the AnnData object from the SRR library metadata provided by Kotliar *et al.^37^*. The cell barcodes were filtered based on a minimum number of UMI counts of 125 obtained from the knee plot of sorted total UMI counts per cell (Extended Data Fig. 3a), resulting in a mean UMI count of 1,401 after filtering. The cells were further filtered based on a maximum percentage of mitochondrial genes of 10 %, based on a combination of macaque and dog mitochondrial genes facilitated by Scanpy^84^ (v1.9.3) and gget^45^ (v0.28.0). Cells were categorized as macaque if a maximum of 10 % of their UMIs originated from dog genes and vice versa (Extended Data Fig. 3b). Macaque and MDCK cells were normalized separately using log(CP10k + 1) with Scanpy’s normalize_total defaults of target sum 10,000 and log1p.

#### Macaque cell clustering and cell type assignment

The macaque gene count matrix was transformed by PCA to 50 dimensions applied using the log-normalized counts filtered for highly variable genes using Scanpy’s highly_variable_genes. Next, we computed nearest neighbors and conducted Leiden clustering^58^ using Scanpy, resulting in 19 Leiden clusters. We found that EVD time points were highly concordant across sequencing libraries, suggesting the lack of a batch effect (Fig. 7a, also see GitHub code repository). Each cluster was manually annotated with a cell type based on the expression of previously established marker genes^37^ (Extended Data Fig. 3d). Cluster ‘Undefined 1’ was omitted because it only contained 12 cells. Gene names and descriptions for Ensembl IDs without annotations were obtained using gget^45^.

#### Virus alignment with different masking options

For each masking option, we quantified the gene expression in each of the 106 datasets from <u>GSE158390</u> using kallisto with the ‘batch’ and ‘batch-barcodes’ arguments to process all files simultaneously while keeping track of each batch and with the ‘x’-string ‘0,0,12:0,12,20:1,0,0’ to match the Seq-Well technology. kallisto translated search was initiated in the ‘kallisto index’ and ‘kallisto bus’ commands by adding the ‘—aa’ flag. Following the alignment to PalmDB with any of the masking options, cell barcodes were corrected using the barcode on-list generated during the alignment to the host as described above.

##### No mask

The raw sequencing reads were aligned to the optimized PalmDB reference files (see ‘Optimization of PalmDB’ above) using kallisto translated search.

##### D-list genome + transcriptome

The raw sequencing reads were aligned to the optimized PalmDB reference using kallisto translated search with the added argument ‘d-list’, which was passed the concatenated macaque genome and transcriptome (Mmul_10 Ensembl version 109), and dog genome and transcriptome (ROS_Cfam_1.0 Ensembl version 109). For D-list masking options including only the genomes or transcriptomes (Extended Data Fig. 4a), only the genome or transcriptome files from both species were concatenated and passed to the ‘d-list’ argument, respectively.

##### D-list genome + transcriptome + ambiguous reads filtered

This workflow was performed as described above for the ‘D-list genome + transcriptome’ with an unreleased version of kallisto where ambiguous reads in the D-list will be thrown out as host instead of being assigned to virus (Extended Data Fig. 4a). We explored this option to investigate the effect of ambiguous reads during D-list masking. However, we found that the alignment results did not notably differ from the masking option ‘D-list genome + transcriptome’ (Fig. 4a and Extended Data Fig. 5).

##### Host read capture with kallisto

The raw sequencing reads were aligned to the combined macaque and dog reference index generated during the alignment to host with ‘kallisto bus’ with the added ‘-n’ flag. The ‘-n’ flag keeps track of the read line number of each aligned read; the line numbers are added to the resulting BUS file. The raw sequencing reads were also aligned to the modified PalmDB with kallisto translated search with the added ‘-n’ flag to obtain all reads that map to viral RdRPs. Subsequently, the BUS file returned by kallisto translated search was split into reads that only aligned to viral RdRPs and reads that also aligned to host based on the read line numbers in the BUS files. This step was performed using ‘bustools capture’ to, first, obtain all reads that belonged to a single batch file (of the 106 dataset files), and, second, capture all reads that also aligned to host.

##### Host read capture with kallisto + D-list genome + transcriptome

Host reads were captured with kallisto as described above under ‘Host read capture with kallisto’. However, during the alignment of the raw sequencing reads to PalmDB with the ‘-n’ flag, we also used the ‘d-list’ flag to mask the host genomes and transcriptomes as described above under ‘D-list genome + transcriptome’.

##### Prior alignment to host with bwa

bwa^54^ v0.7.17 was installed from source. The ‘bwa index’ command was used to generate a bwa index from the concatenated macaque and dog genomes (Mmul_10 and ROS_Cfam_1.0 from Ensembl v109). The raw sequencing reads were subsequently aligned to the bwa index using the ‘bwa mem’ command, aligning each file separately. For each FASTQ file, the names of all unmapped reads were extracted using ‘samtools view’ (SAMtools^40^ v1.6), and a new FASTQ file including only unmapped sequences was generated using the ‘seqtk subseq’ command (v1.4; https://github.com/lh3/seqtk). The resulting FASTQ files containing the sequencing reads that did not map to the host genomes were aligned to the optimized PalmDB reference files using kallisto translated search.

#### Extraction and BLAST alignment of viral reads

Randomly selected sequencing reads from three libraries that included reads that mapped to the viruses of interest were aligned to the optimized PalmDB with kallisto translated search including the ‘-n’ flag, without any host read masking. Reads that mapped to the viruses of interest were subsequently captured and extracted from the raw sequencing FASTQ files using ‘bustools capture’ and ‘bustools extract’.

BLAST+^56^ v2.14.1 was installed from source and the BLAST nt database was downloaded using the update_blastdb.pl command. 10 reads were randomly chosen for each target virus for each library and were BLASTed against the nt database using the blastn algorithm. Sequences that aligned to the polyA tail were recognized by the occurrence of ‘AAAAAAAAAAAA’ or ‘TTTTTTTTTTTT’ in the aligned part of the subject or query sequences and removed from the results. BLAST results were subsequently plotted using pyCirclize.Circos (v1.0.0; https://github.com/moshi4/pyCirclize).

#### Virus quality control

The viral count matrix generated using the ‘Host read capture with kallisto + D-list genome + transcriptome’ masking workflow was converted to h5ad using kb_python.utils.kb_utils and read into Python using anndata v0.8.0. Metadata such as donor animal, the presence of an MDCK spike-in, and time point were added to the AnnData object from the SRR library metadata provided by Kotliar *et al.^37^*. For each cell, the host species and cell type were added from the host matrices generated as described above. The virus count matrix was subsequently binarized, such that for each cell, each virus was either present or absent. The viruses were thresholded to viruses that were observed in ≥ 0.05 % of cells in either species.

#### Virus categorization into shared, ‘macaque only’, and ‘MDCK only’ viruses

For each virus ID, the virus was defined as ‘shared’ if the fold change between the fraction of positive macaque cells and the fraction of positive MDCK cells was less than or equal to 2. Viruses were assigned the category ‘macaque only’ if the virus was seen in ≥ 0.05 % of macaque cells and ≤ 7 MDCK cells, and vice versa for the category ‘MDCK only’. These thresholds were defined based on the percentages of positive cells observed for each virus in each species, as shown in Fig. 6b.

#### Generation of the Krona plot

KronaTools^62^ v2.8.1 was installed from source. We generated a data frame containing the total numbers of positive cells for each sOTU seen in ≥ 0.05 % of macaque cells for each animal and time point (including only cells that passed host cell quality control). The ktImportText tool was used to generate a Krona plot HTML file from a text file generated from this data frame.

#### Logistic regression models to predict viral presence based on host gene expression

Logistic regression models the log odds of an event as a weighted linear combination of some predictor variables. That is, the natural log of the ratio of the probability *p* that an event occurs to the probability that it does not occur is modeled:

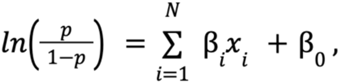

where each *x_i_* is a predictor variable with corresponding weight *β*_i_ and *β*_0_ is an intercept. Here, *p* is the probability of viral presence or absence in a given cell, predicted based on a linear combination of normalized host gene count values (denoted as *x* with a total of *G* modeled genes). Viral presence or absence is modeled for a single virus at a time. To control for covariates, we also included animal identifier (denoted as *y* with a total of *A* animals) and time point (denoted as *z* with a total of *T* time points), which were one-hot encoded for fits:

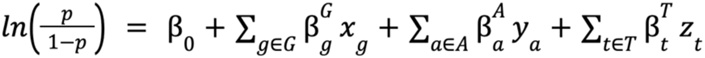

The magnitude of the weight value for each predictor variable corresponds to that variable’s influence on event probability, with large positive weights increasing the probability and large negative weights decreasing the probability of the event. Thus, for our purposes, an analysis of gene weights suggests which genes are likely to correlate with viral infection. For models parameterized by highly variable (HV) genes, the host (macaque) matrix was subset to highly variable genes as defined above. To reduce the occurrence of false negative viral counts, the logistic regression models were trained using the viral count matrix obtained without any masking of the host genes. However, the models were trained for viruses that were filtered based on the more conservative masking options (‘macaque only’ and ‘shared’ viruses). To further reduce the occurrence of false negative viral counts, we filtered the virus and host matrices to include only the top 50 % of cells according to the sum of raw host reads per cell before training the models. This was done to reduce the effects introduced by varying sequencing depths. For example, cells with a lower sequencing depth will have a higher likelihood of a false negative viral count.

For viruses with more virus-negative than virus-positive cells, half of the virus-positive cells and an equal number of virus-negative cells were randomly selected to train the logistic regression models. For viruses with more virus-positive than virus-negative cells, half of the virus-negative cells and an equal number of virus-positive cells were randomly selected for training. In both cases, the remaining cells were held out for testing the performance of trained models. Given the cell type specificity of the viruses whose presence could be predicted with high accuracy, we wanted to confirm that we were not simply predicting cell type. To this end, virus-negative training cells were selected to be of the same cell types as virus-positive cells (Extended Data Fig. 8d). The number of training and testing cells for each virus are shown in Extended Data Fig. 8c.

For models that included covariates, donor animal and EVD time point were one-hot encoded and appended to the gene expression training matrix. All models included an intercept. Models were trained with L2 weight regularization using the sklearn.linear_model.LogisticRegression (sklearn v1.0.1) classifier with a maximum of 100 iterations to predict the probability of viral presence at single-cell resolution. Virus-positive cells were assigned class label 1, and virus-negative cells were assigned class label 0. All four possible combinations of two modeling choices (highly variable versus all genes, and covariates versus no covariates) were tested, and the results are shown in Fig. 8c. Accuracy, specificity, and sensitivity were calculated for each model on the held-out testing cells (Extended Data Fig. 8a). A negative control where labels of viral presence and absence for each virus were randomly scrambled in the training data was included in the modeling experiments. For the scrambled labels, the original ratio of virus-positive to virus-negative cells was maintained.

All results were averaged across models generated using six different random seeds for parameter optimization and random selection of cells for training and testing.

#### Enrichment analysis of predictive genes

Of the top 50 highly variable macaque genes with the largest positive average weights in the regression model we selected those for which the standard deviation of the weights was less than half than the lowest weight. Here, we used the model trained on highly variable genes with covariates donor animal and time point. The gene weight distributions and thresholds are shown in Extended Data Fig. 9a. Approximately one third of the macaque Ensembl IDs did not have annotated gene names. We used gget^45^ to translate annotated Ensembl IDs to gene symbols and to perform enrichment analysis on the returned gene symbols using Enrichr^71^ against the 2023 Gene Ontology (GO) Biological Processes database (‘GO_Biological_Process_2023’)^72^. The reported P values were corrected with the Benjamini-Hochberg method^73^.

## Data availability

**Table 1:** Availability of data analyzed in this paper.

| Sample | Methods | Data shown in | DOI | GEO accession |
| --- | --- | --- | --- | --- |
| Lung autopsy samples from COVID-19 patients | Bulk RNA sequencing; RNA-ISH | Fig. 3a | https://doi.org/10.1038/s41467-020-20139-7 | [GSE150316](https://www.ncbi.nlm.nih.gov/geo/accession/GSE150316) |
| Self-collected saliva, anterior nares swab, and oropharyngeal swab samples from individuals enrolled in a COVID-19 household transmission study | Bulk RNA sequencing with viral surveillance panel enrichment (Illumina Cat. 20040536 and 20088154); RT-qPCR | Fig. 3a | https://doi.org/10.1128/spectrum.03873-22<br>https://doi.org/10.1093/pnasnexus/pgad033 | Raw sequencing data is not publicly available per participant privacy practices |
| SARS-CoV-2 infected human iPSC derived cardiomyocytes | SMART-Seq V4 | Fig. 3a | https://doi.org/10.1016/j.xcrm.2020.100052 | [GSM4548303](https://www.ncbi.nlm.nih.gov/geo/accession/GSM4548303) |
| Blood samples from rhesus macaques infected with *Zaire ebolavirus* | Seq-Well S<sup>3</sup>; RT-qPCR | Fig. 2-8; Extended Data Fig. 1-9 | https://doi.org/10.1016/j.cell.2020.10.002 | [GSE158390](https://www.ncbi.nlm.nih.gov/geo/accession/GSE158390) |
| Lungs from APOE knock-in mice infected with SARS-CoV-2 | SPLiT-Seq (Parse Biosciences) | Extended Data Fig. 4c | https://doi.org/10.1038/s41586-022-05344-2 | [GSM5974202](https://www.ncbi.nlm.nih.gov/geo/accession/GSM5974202) |

**Table 2:** Availability of data generated in this paper. The data is available on Caltech Data under the DOIs 10.22002/krqmp-5hy81 and 10.22002/k7xqw-88d74.

| File | Description | Category |
| --- | --- | --- |
| viral sequences in laboratory reagents.h5ad | Count matrix containing virus-like sequences found in sequencing libraries comprised of only sterile water and laboratory reagents | Alignment of 'blank' sequencing libraries to the PalmDB |
| host alignment results.zip | Raw alignment results obtained by kallisto after alignment to the macaque and dog (to account for the MDCK spike-in) transcriptomes | Alignment of the macaque PBMC data<sup>37</sup> to the host transcriptome(s) |
| host QC.h5ad | Filtered count matrix containing all host cells |  |
| canis QC norm leiden.h5ad | Filtered and clustered count matrix containing MDCK cells |  |
| macaque QC norm leiden.h5ad | Filtered and clustered count matrix containing macaque cells |  |
| macaque QC norm leiden celltypes.h5ad | Filtered and clustered count matrix containing macaque cells with cell type assignments |  |
| virus no mask alignment results.zip | Raw alignment results obtained by kallisto translated search after alignment to the PalmDB without masking host sequences | Alignment of the macaque PBMC data<sup>37</sup> to the PalmDB for the detection of viral RNA with different workflows for the masking of host genome(s) and transcriptome(s) |
| virus no mask.h5ad | Count matrix obtained through the alignment above with added metadata |  |
| virus dlist cdna alignment results.zip | Raw alignment results obtained by kallisto translated search after alignment to the PalmDB while masking host transcriptome(s) using the D-list |  |
| virus dlist cdna.h5ad | Count matrix obtained through the alignment above with added metadata |  |
| virus dlist dna alignment results.zip | Raw alignment results obtained by kallisto translated search after alignment to the PalmDB while masking host genome(s) using the D-list |  |
| virus dlist dna.h5ad | Count matrix obtained through the alignment above with added metadata |  |
| virus dlist cdna dna alignment results.zip | Raw alignment results obtained by kallisto translated search after alignment to the PalmDB while masking host genome(s) and transcriptome(s) using the D-list |  |
| virus dlist cdna dna.h5ad | Count matrix obtained through the alignment above with added metadata |  |
| virus dlist cdna dna amb alignment results.zip | Raw alignment results obtained by kallisto translated search after alignment to the PalmDB while masking host genome(s) and transcriptome(s) using the D-list + forcing ambiguous sequences to be discarded |  |
| virus dlist cdna dna ambiguous.h5ad | Count matrix obtained through the alignment above with added metadata |  |
| virus host capture alignment results.tar.gz | Raw alignment results obtained by kallisto translated search after alignment to the PalmDB + reads that align to the host transcriptome(s) were captured |  |
| virus host-captured.h5ad | Count matrix obtained through the alignment above with added metadata |  |
| virus host capture dlist cdna dna alignment results.tar.gz | Raw alignment results obtained by kallisto translated search after alignment to the PalmDB while masking host genome(s) and transcriptome(s) using the D-list + reads that align to the host transcriptome(s) were captured |  |
| virus host-captured dlist cdna dna.h5ad | Count matrix obtained through the alignment above with added metadata |  |
| bwa unmapped reads.tar.gz | Raw sequencing files obtained after removal of host sequences based on alignment with bwa |  |
| virus bwa alignment results.zip | Raw alignment results obtained by kallisto translated search after alignment to the PalmDB after reads that align to the host genome(s) with bwa were removed |  |
| virus bwa.h5ad | Count matrix obtained through the alignment above with added metadata |  |
| models.zip | Logistic regression models to predict viral presence based on host gene expression | Logistic regression models |
| palmdb human dlist cdna dna.idx | Pre-computed PalmDB reference index with human genomic and transcriptomic sequences masked using D-list | Pre-computed references for future use with kallisto translated search |
| palmdb mouse dlist cdna dna.idx | Pre-computed PalmDB reference index with mouse genomic and transcriptomic sequences masked using D-list |  |

The PalmDB reference files optimized for use with kallisto translated search for the identification of viral sequences in bulk and single-cell RNA sequencing data are available here: https://tinyurl.com/4wd33rey.

The data generated in this paper is freely and publicly available on Caltech Data under the DOIs 10.22002/krqmp-5hy81 and 10.22002/k7xqw-88d74.

## Code availability

The code used to generate all of the results and figures reported in this paper, starting from the raw sequencing reads, can be found here: https://github.com/pachterlab/LSCHWCP_2023. The code is organized by figure panel and provided in immediately executable Google Colab notebooks to maximize the reproducibility of the results and methods described in this manuscript.

## Acknowledgments

We would like to thank Astrid Bryon for her advice and helpful discussions on the applicability of kallisto translated search for the detection of viral RNA. We would like to thank Aaron Lin, Lisa Hensley, Dylan Kotliar, and Pardis Sabeti for helping us understand and work with their data. We would like to thank Christina Middle, Morgan Roos, and Scott Kuersten from Illumina for generating the bulk RNA sequencing data included in Fig. 3a (second panel). The illustrations in Figures 1, 4, and 6 were created with BioRender.com.

## Ethics declarations

Competing Interests: LL, DKS, and LP are listed as inventors of a patent application relating to the work. The patent application was submitted through the Technology Transfer Office of the California Institute of Technology (Caltech), with Caltech being the patent applicant.

## Extended Data

**Extended Data Table 1:**
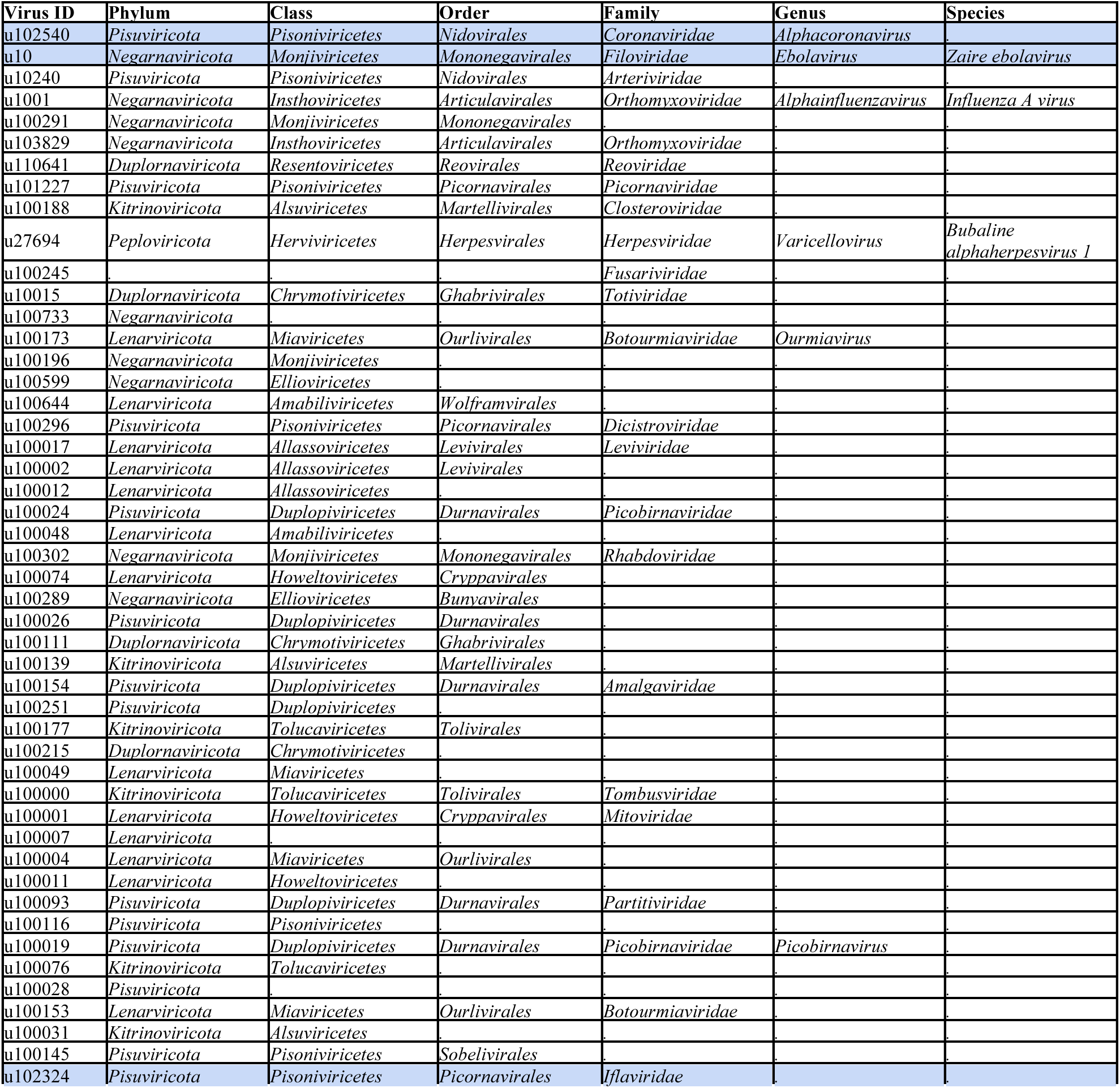
Virus ID to species-like operational taxonomic unit (sOTU) mapping for the most highly expressed viruses (in the same order as shown in *Fig. 6d*). Virus IDs that are further mentioned in the paper are marked in blue. Virus IDs not included in this list are of unknown taxonomy across all taxonomic ranks.

| Virus ID | Phylum | Class | Order | Family | Genus | Species |
| --- | --- | --- | --- | --- | --- | --- |
| u102540 | Pisuviricota | Pisoniviricetes | Nidovirales | Coronaviridae | Alphacoronavirus |  |
| u10 | Negarnaviricota | Monjiviricetes | Mononegavirales | Filoviridae | Ebolavirus | Zaire ebolavirus |
| u10240 | Pisuviricota | Pisoniviricetes | Nidovirales | Arteriviridae |  |  |
| u1001 | Negarnaviricota | Insthoviricetes | Articulavirales | Orthomyxoviridae | Alphainfluenzavirus | Influenza A virus |
| u100291 | Negarnaviricota | Monjiviricetes | Mononegavirales |  |  |  |
| u103829 | Negarnaviricota | Insthoviricetes | Articulavirales | Orthomyxoviridae |  |  |
| u110641 | Duplornaviricota | Resentoviricetes | Reovirales | Reoviridae |  |  |
| u101227 | Pisuviricota | Pisoniviricetes | Picornavirales | Picornaviridae |  |  |
| u100188 | Kitrinoviricota | Alsuviricetes | Martellivirales | Closteroviridae |  |  |
| u27694 | Peploviricota | Herviviricetes | Herpesvirales | Herpesviridae | Varicellovirus | Bubaline alphaherpesvirus 1 |
| u100245 |  |  |  | Fusariviridae |  |  |
| u10015 | Duplornaviricota | Chrymotiviricetes | Ghabrivirales | Totiviridae |  |  |
| u100733 | Negarnaviricota |  |  |  |  |  |
| u100173 | Lenarviricota | Miaviricetes | Ourlivirales | Botourmiaviridae | Ourmiavirus |  |
| u100196 | Negarnaviricota | Monjiviricetes |  |  |  |  |
| u100599 | Negarnaviricota | Ellioviricetes |  |  |  |  |
| u100644 | Lenarviricota | Amabiliviricetes | Wolframvirales |  |  |  |
| u100296 | Pisuviricota | Pisoniviricetes | Picornavirales | Dicistroviridae |  |  |
| u100017 | Lenarviricota | Allasoviricetes | Levivirales | Leviviridae |  |  |
| u100002 | Lenarviricota | Allasoviricetes | Levivirales |  |  |  |
| u100012 | Lenarviricota | Allasoviricetes |  |  |  |  |
| u100024 | Pisuviricota | Duplopiviricetes | Durnavirales | Picobirnaviridae |  |  |
| u100048 | Lenarviricota | Amabiliviricetes |  |  |  |  |
| u100302 | Negarnaviricota | Monjiviricetes | Mononegavirales | Rhabdoviridae |  |  |
| u100074 | Lenarviricota | Howeltoviricetes | Cryppavirales |  |  |  |
| u100289 | Negarnaviricota | Ellioviricetes | Bunyavirales |  |  |  |
| u100026 | Pisuviricota | Duplopiviricetes | Durnavirales |  |  |  |
| u100111 | Duplornaviricota | Chrymotiviricetes | Ghabrivirales |  |  |  |
| u100139 | Kitrinoviricota | Alsuviricetes | Martellivirales |  |  |  |
| u100154 | Pisuviricota | Duplopiviricetes | Durnavirales | Amalgaviridae |  |  |
| u100251 | Pisuviricota | Duplopiviricetes |  |  |  |  |
| u100177 | Kitrinoviricota | Tolucaviricetes | Tolivirales |  |  |  |
| u100215 | Duplornaviricota | Chrymotiviricetes |  |  |  |  |
| u100049 | Lenarviricota | Miaviricetes |  |  |  |  |
| u100000 | Kitrinoviricota | Tolucaviricetes | Tolivirales | Tombusviridae |  |  |
| u100001 | Lenarviricota | Howeltoviricetes | Cryppavirales | Mitoviridae |  |  |
| u100007 | Lenarviricota |  |  |  |  |  |
| u100004 | Lenarviricota | Miaviricetes | Ourlivirales |  |  |  |
| u100011 | Lenarviricota | Howeltoviricetes |  |  |  |  |
| u100093 | Pisuviricota | Duplopiviricetes | Durnavirales | Partitiviridae |  |  |
| u100116 | Pisuviricota | Pisoniviricetes |  |  |  |  |
| u100019 | Pisuviricota | Duplopiviricetes | Durnavirales | Picobirnaviridae | Picobirnavirus |  |
| u100076 | Kitrinoviricota | Tolucaviricetes |  |  |  |  |
| u100028 | Pisuviricota |  |  |  |  |  |
| u100153 | Lenarviricota | Miaviricetes | Ourlivirales | Botourmiaviridae |  |  |
| u100031 | Kitrinoviricota | Alsuviricetes |  |  |  |  |
| u100145 | Pisuviricota | Pisoniviricetes | Sobelivirales |  |  |  |
| u102324 | Pisuviricota | Pisoniviricetes | Picornavirales | Iflaviridae |  |  |

**Extended Data Fig. 1:**
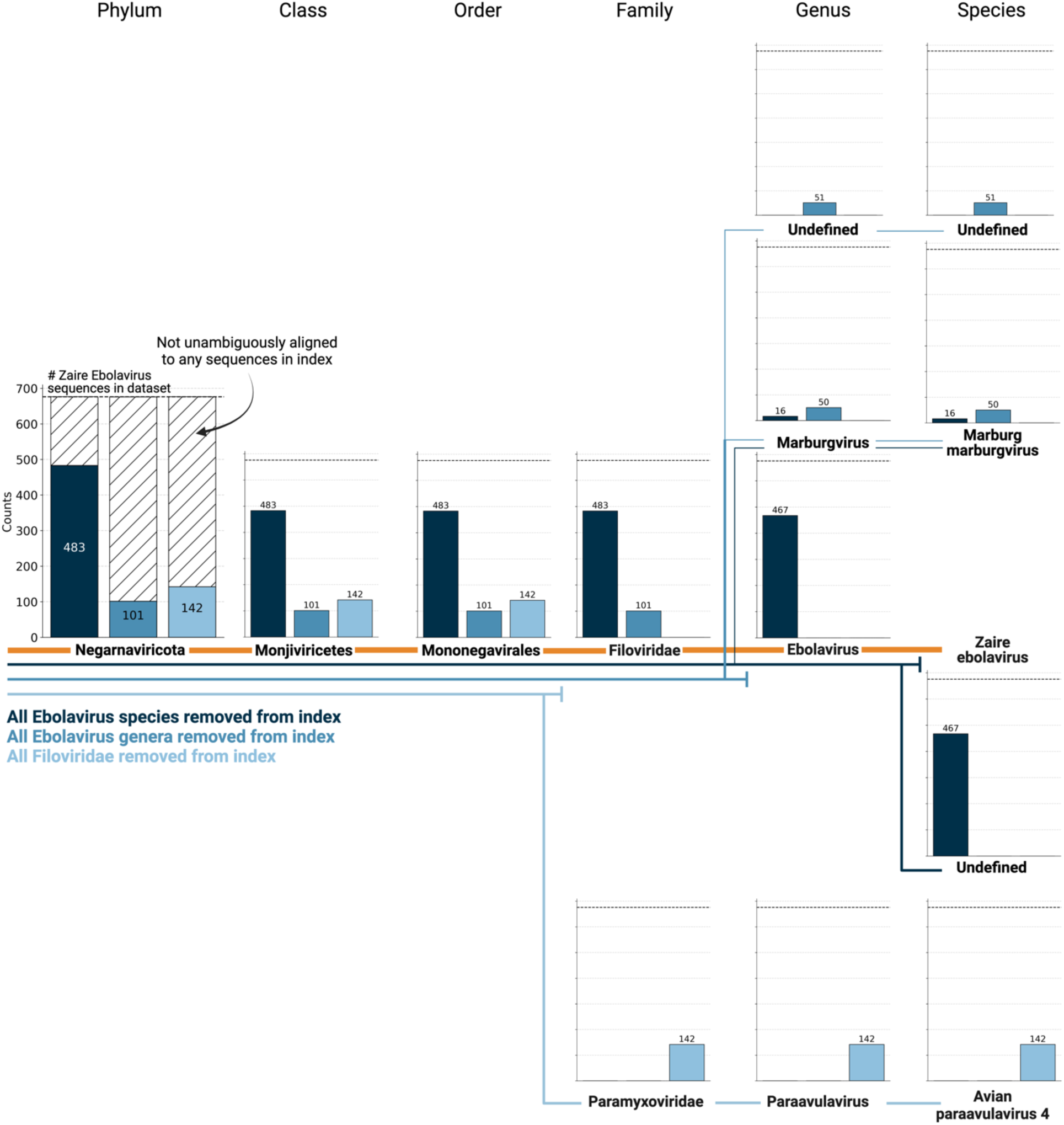
676 ZEBOV RdRP sequences were identified by aligning a subset of 100,000,000 single-cell RNA sequencing reads of macaque PBMC samples obtained at 8 days post-infection with ZEBOV^37^ to the optimized PalmDB using kallisto translated search. We subsequently aligned the sequences to PalmDB reference indices from which (i) all Ebolavirus species were removed (dark blue), (ii) all Ebolavirus genera were removed (medium blue), or (iii) all Filoviridae were removed (light blue). In each scenario, a subset of sequences aligned to the nearest remaining relative based on the main taxonomic rank, suggesting that kallisto translated search can detect the highly conserved RdRP of a large number of viral species, beyond the number of sequences in the PalmDB database, while still providing reliable sOTU-based taxonomic assignment of lower-rank taxonomies.

**Extended Data Fig. 2:**
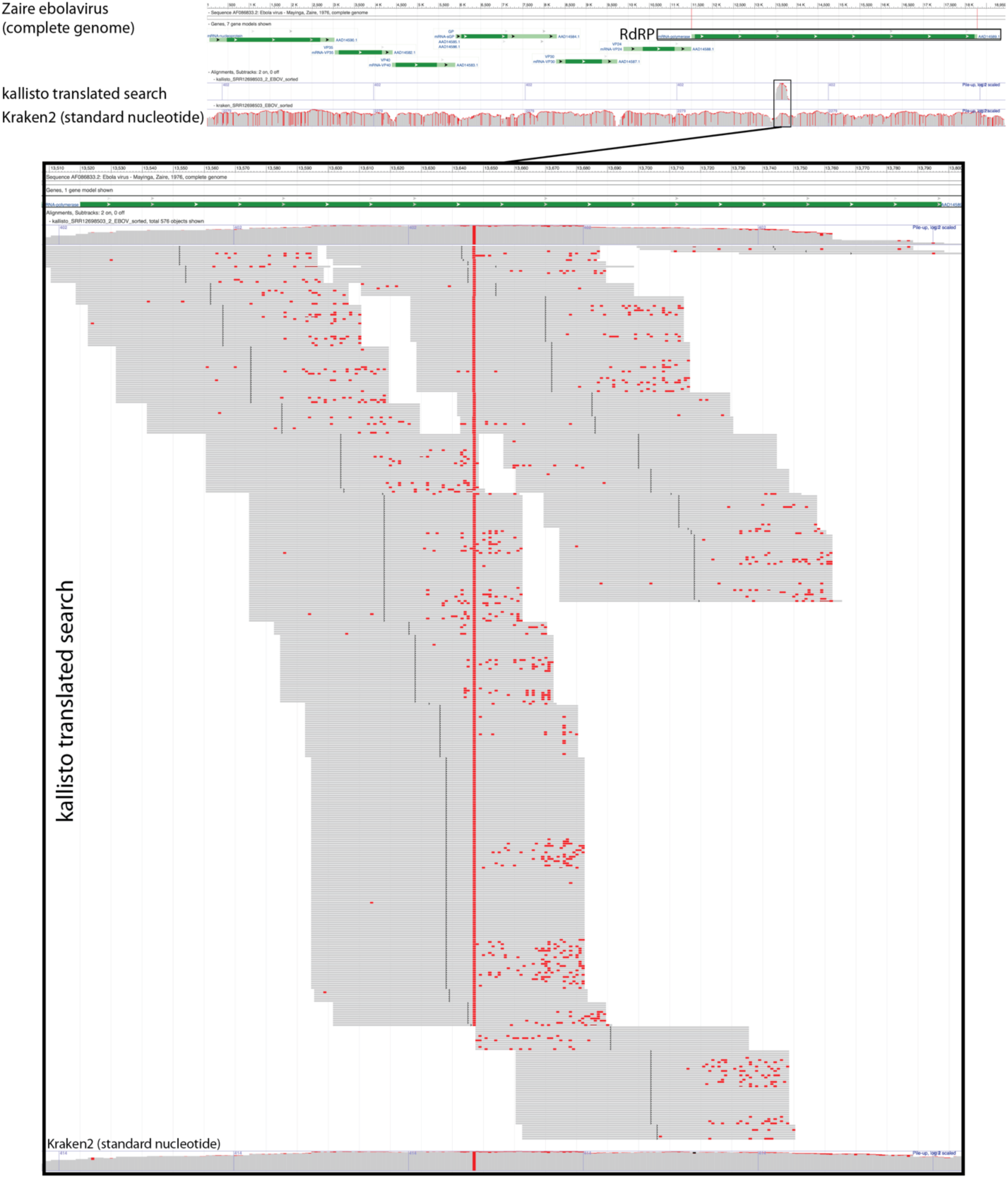
Visualization of the identification of RdRP sequences with kallisto translated search. We selected a subset of 100,000,000 reads obtained using Seq-Well sequencing of macaque peripheral blood mononuclear cell (PBMC) samples obtained at 8 days post-infection with ZEBOV^37^. We aligned the reads to the PalmDB amino acid sequences with kallisto translated search. We also aligned the reads to the complete ZEBOV nucleotide genome using Kraken2 (standard nucleotide alignment)^27^. Aligned reads from both alignments were extracted and realigned to the ZEBOV genome using bowtie2^39^, a BAM file was created using SAMtools^40^, and the alignment was subsequently visualized NCBI Genome Workbench^41^.

**Extended Data Fig. 3:**
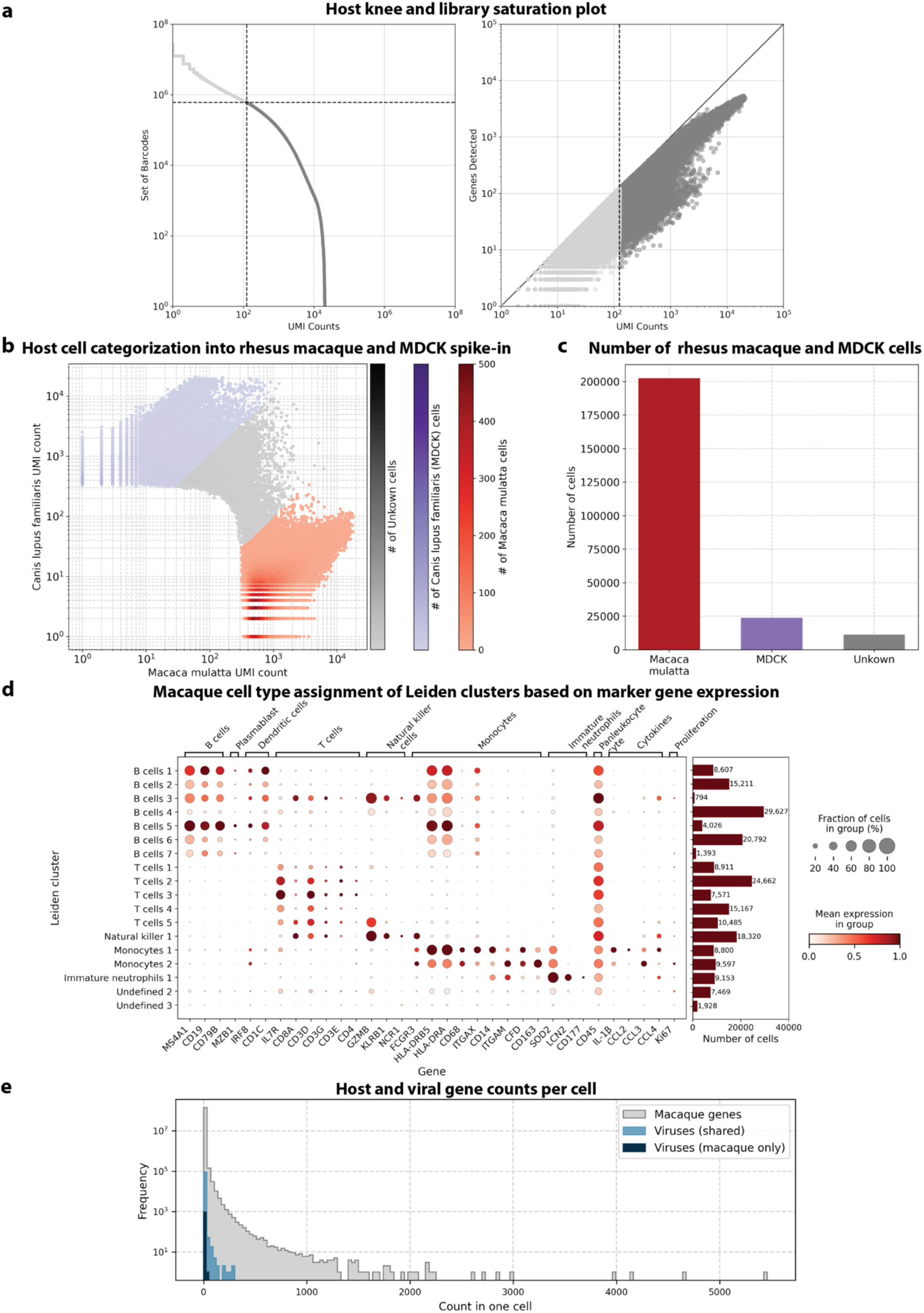
**a**, Knee plot of sorted total UMI counts per cell and library saturation plot of host (rhesus macaque and MDCK) cells sequenced by Kotliar et al.^37^ **b,** Canis lupus (dog/MDCK) over Macaca mulatta (macaque) UMI count for each cell. Cells were categorized as macaque if a maximum of 10 % of their UMIs originated from dog genes and vice versa. **c**, The obtained numbers of macaque, dog (MDCK), and uncategorized cells after species separation. **d**, Mean expression of marker genes used for cell type assignment per macaque Leiden cluster. The barplot shows the number of cells in each cluster. Cluster ‘Undefined 1’ was omitted because it only contained 12 cells. **e**, Frequency of host and viral gene counts in individual cells.

**Extended Data Fig. 4:**
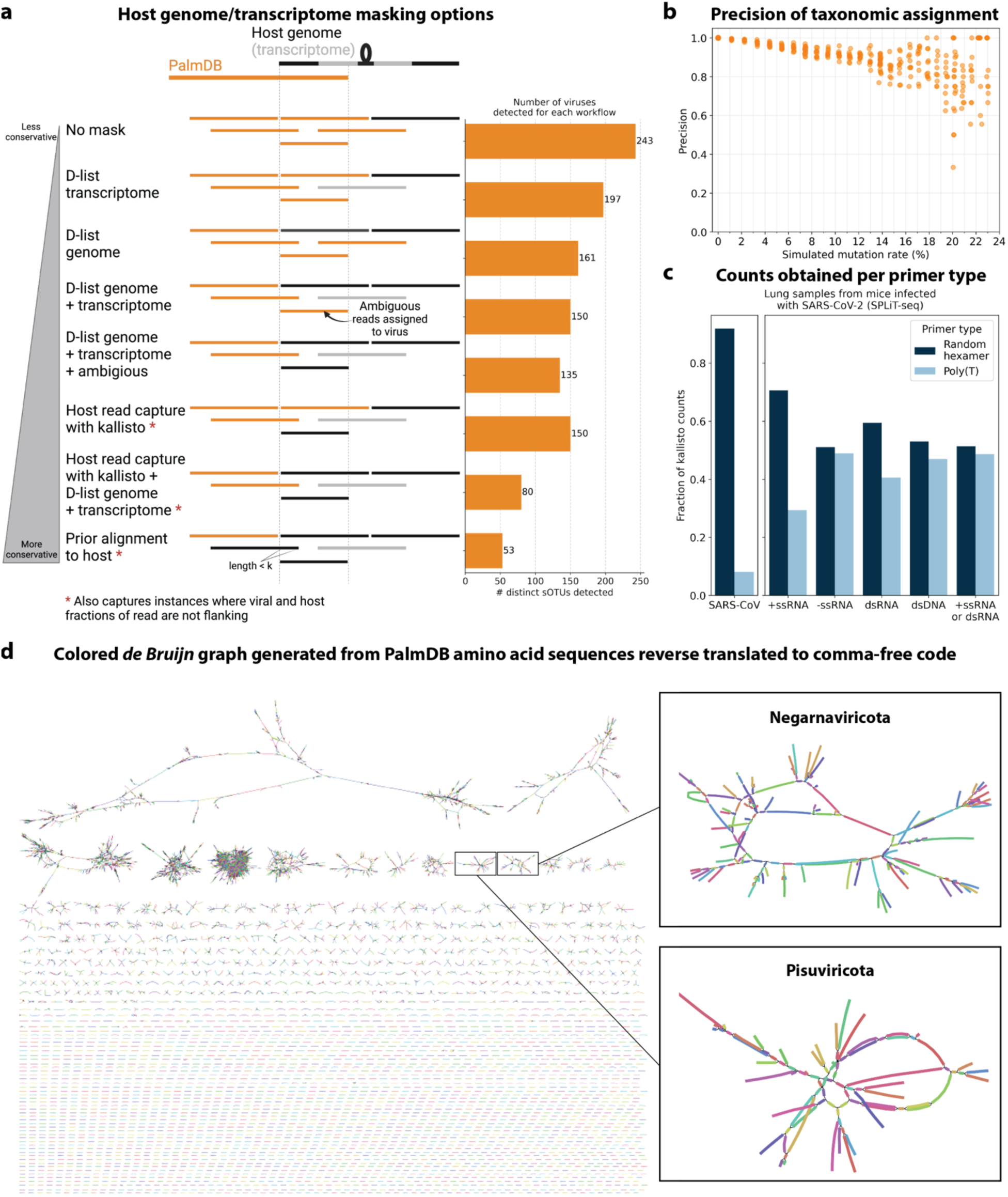
**a**, Schematic overview of different host masking options, extending the masking options shown in *Fig. 4a*. Reads that align to PalmDB and are considered viral are marked in orange, and reads that align to the host genome or transcriptome are marked in black or grey, respectively. The barplot shows the number of distinct sOTUs, defined by distinct virus IDs, observed in ≥ 0.05 % of cells for each workflow. **b**, Precision of species-level taxonomic assignment at increasing simulated mutation rates. Mutation-Simulator^48^ was used to add random single nucleotide base substitutions to 676 ZEBOV RdRP sequences obtained by Seq-Well sequencing^37^ at increasing mutation rates. We performed 10 simulations per mutation rate. The sequences were subsequently aligned using kallisto translated search against the complete PalmDB. The recall percentages at each mutation rate are shown in *Fig. 2c*. **c**, Fraction of counts obtained for the known viral infection (here, SARS-CoV-2) and per viral strandedness of other sOTUs per primer type. Lung samples from mice infected with SARS-CoV2 were sequenced with SPLiT-Seq^85^ and aligned to PalmDB using kallisto translated search using the D-list to mask the host (here, mouse) genome. The plot shows the fraction of counts obtained for SARS-CoV as well as all sOTUs of different strandedness per primer type. **d**, The de Bruijn graph generated from the reverse translated PalmDB sequences in the kallisto translated search workflow, visualized and colored using Bandage v0.8.1^86^.

**Extended Data Fig. 5:**
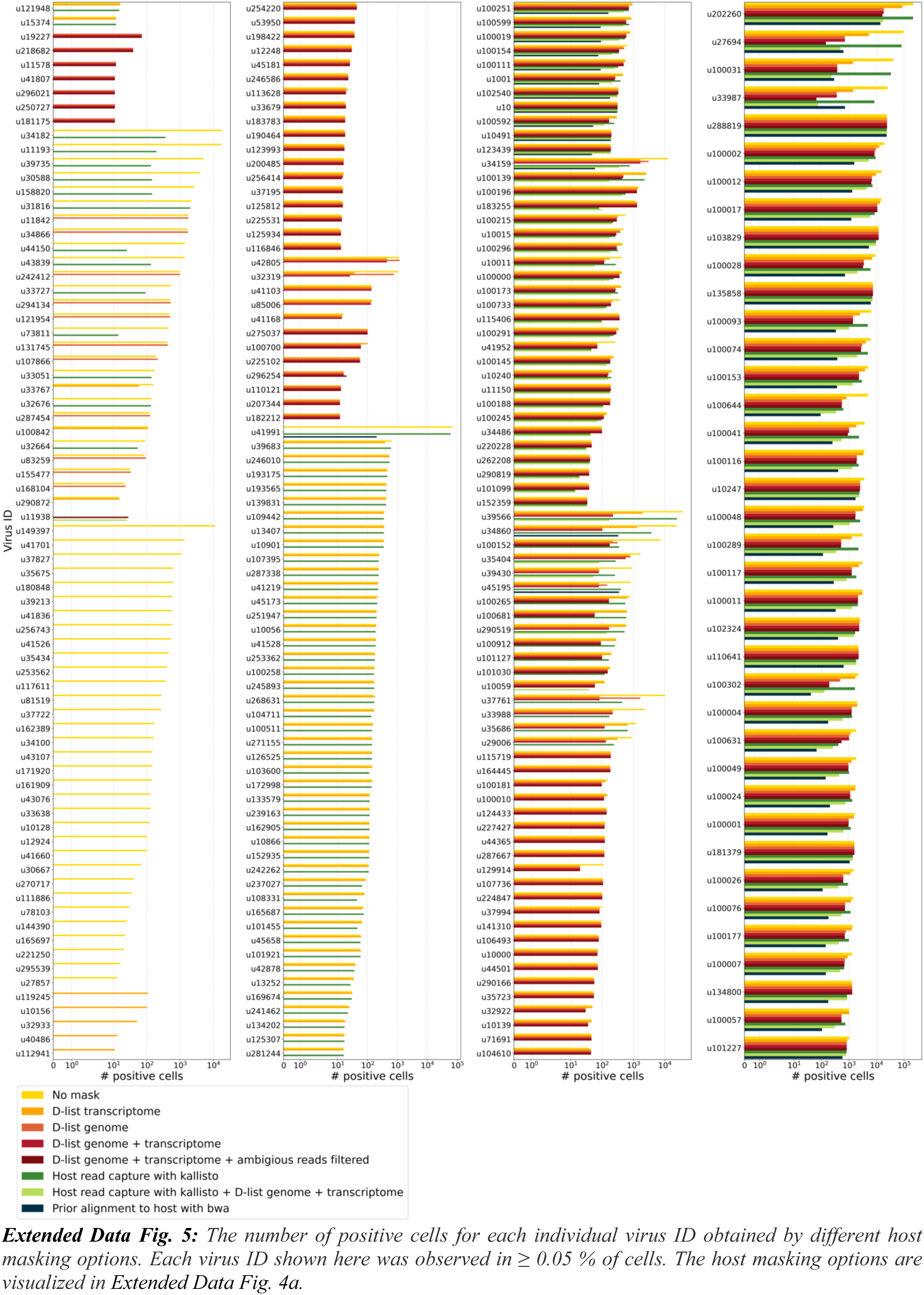
The number of positive cells for each individual virus ID obtained by different host masking options. Each virus ID shown here was observed in ≥ 0.05 % of cells. The host masking options are visualized in Extended Data Fig. 4a.

**Extended Data Fig. 6:**
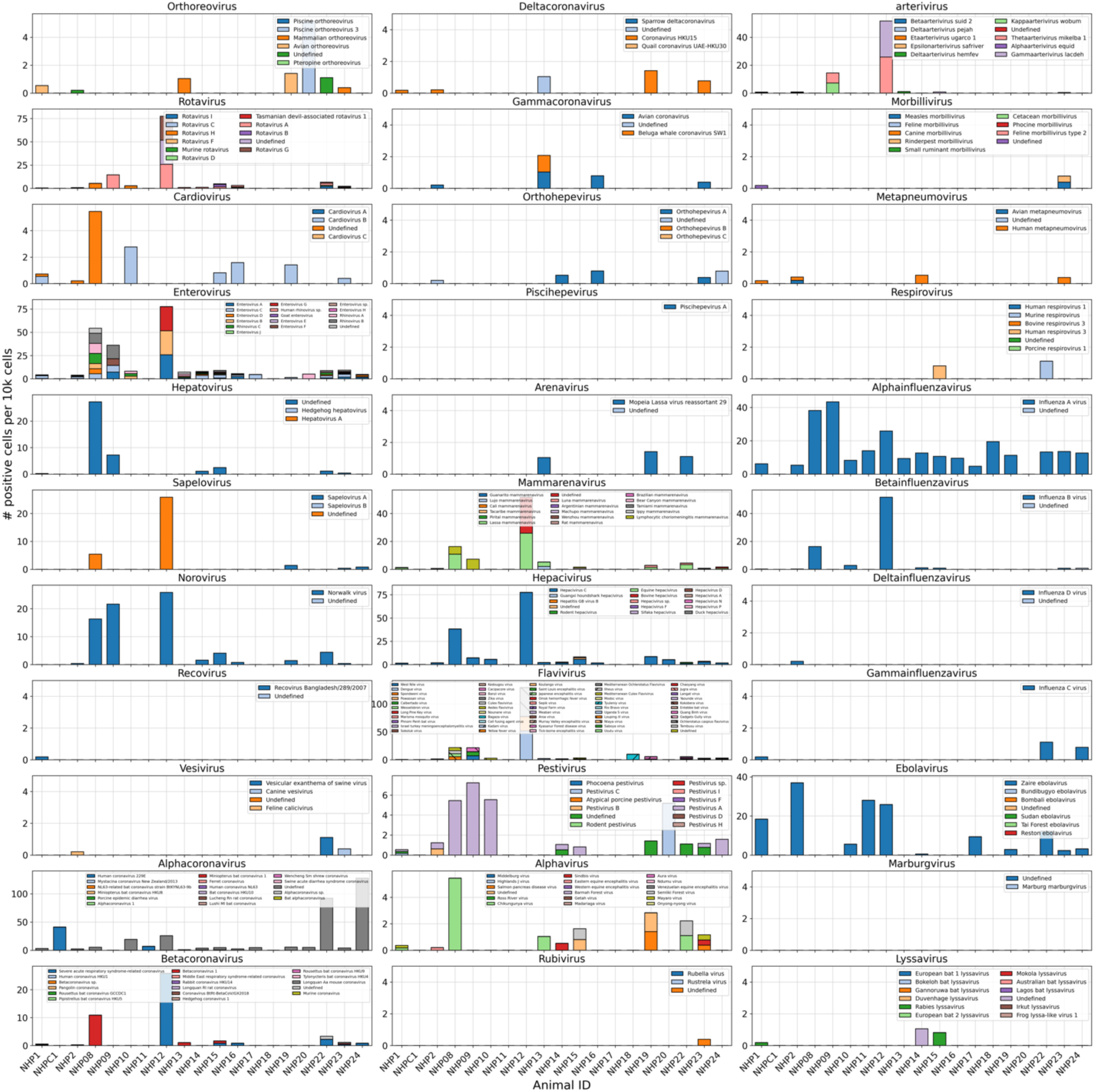
Number of positive cells per 10k cells for virus-like sequences from genera known to infect rhesus macaques^61^ in the data from Kotliar et al.^37^ analyzed using kallisto translated search with PalmDB. Host sequences were masked using the D-list option with the host genomes and transcriptomes, followed by host read capture using kallisto. No quality control thresholding of virus-like sequences was performed prior to generating this plot and the majority of these virus-like sequences were filtered out during quality control, and identification of contaminating sequences.

**Extended Data Fig. 7:**
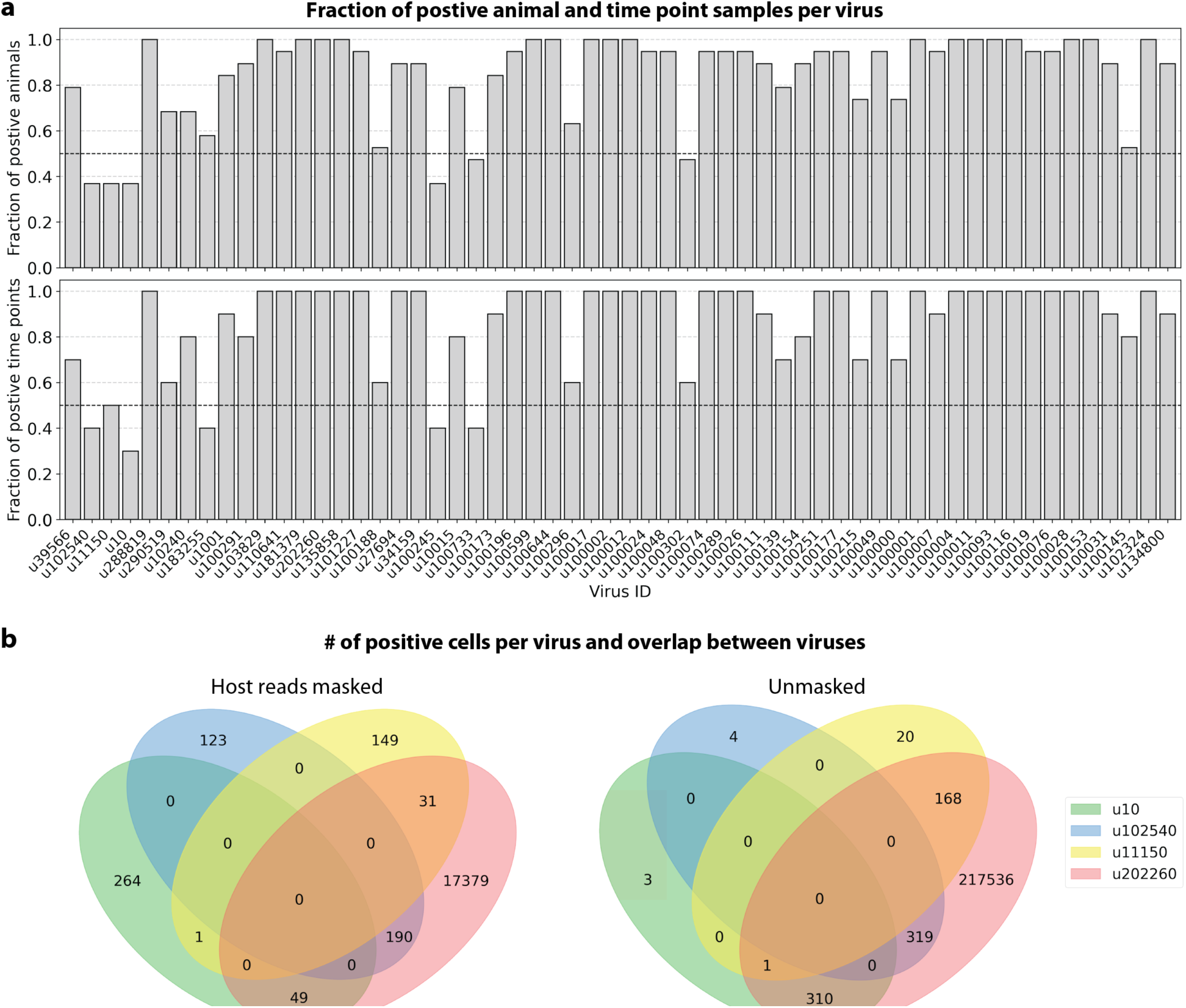
**a**, For each virus ID, the fraction of positive animal (top) and time point (bottom) samples was plotted. A sample was considered positive if at least 0.05 % of cells were positive. **b**, The number of positive cells for each virus ID or any combination of virus IDs for the count matrices generated from host-masked reads (D-list host genome and transcriptome + host transcriptome read capture) (left) and reads without any host masking (right). A large amount of reads for u202260 were masked when conservatively removing host reads (*Fig. 5a*). The plots were generated using PyVenn (https://github.com/tctianchi/pyvenn).

**Extended Data Fig. 8:**
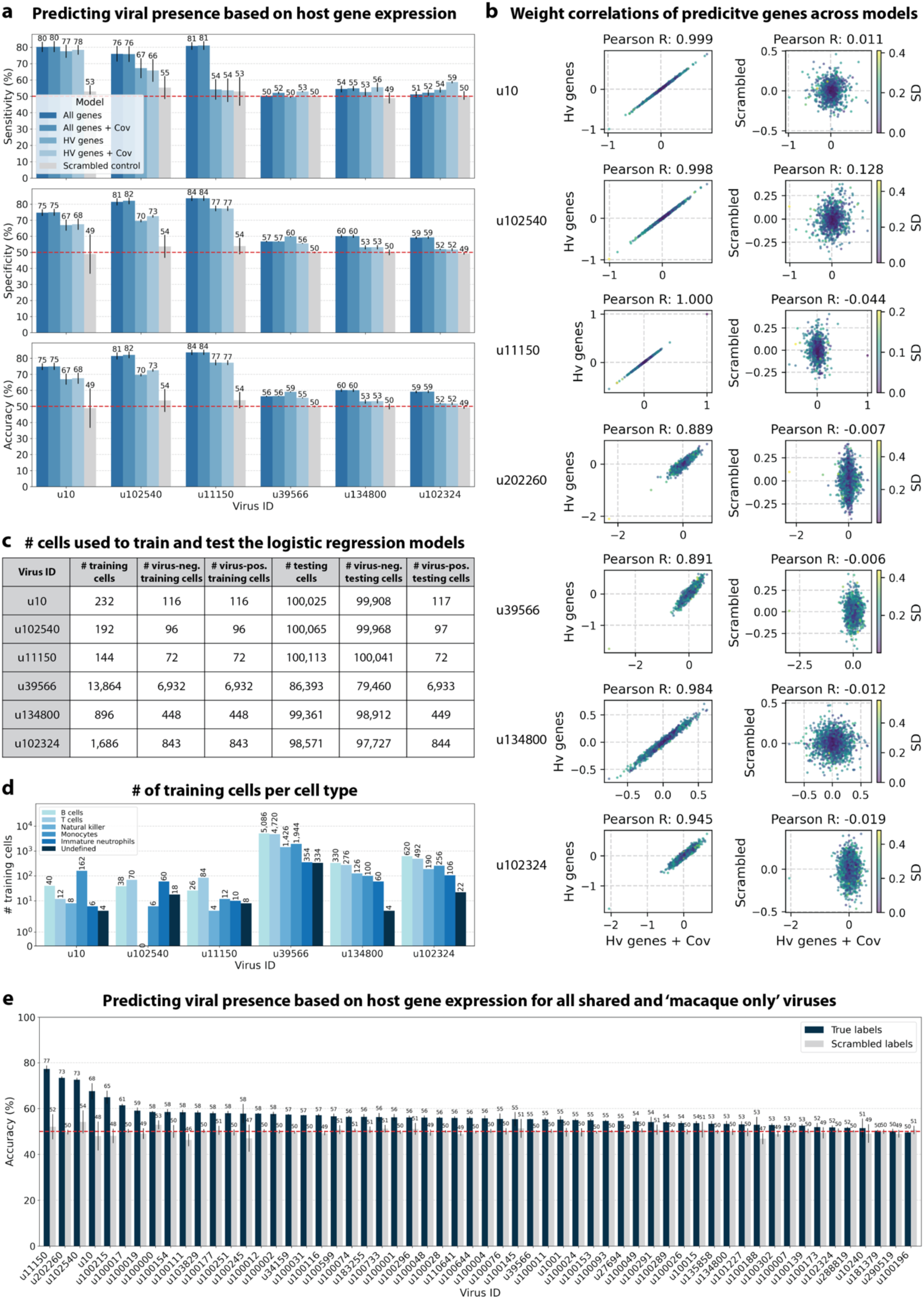
**a**, We trained logistic regression models to predict the presence of specific viruses based on host gene expression at single-cell resolution. The average accuracy, specificity, and sensitivity of the logistic regression models trained on highly variable (HV) or all macaque genes with or without donor animal and EVD time point as covariates are shown for ZEBOV (u10) and five novel virus-like sequences. Error bars indicate the standard deviation between models initialized with different random seeds. As a negative control, viral presence and absence labels were scrambled at random in the training data. **b**, Correlations of the average weights of predictive genes for models trained on HV genes with and without covariates on the real and scrambled labels. The weight correlations are lost when the model is trained using the scrambled labels. Virus IDs with high cell type specificity have slightly higher correlations than those with low cell type specificity. The color bar indicates the standard deviation (SD) of gene weights generated using different random seeds in the model trained on HV genes with covariates. The weights were max normalized between random seeds before computing the average and SD. **c**, The number of cells used to train and test the logistic regression models for ZEBOV (u10) and five novel virus-like sequences. **d**, Total number of training cells per cell type. The total consists of an equal number of virus-positive and -negative cells. **e**, Average prediction accuracy of models trained on HV genes with donor animal and EVD time point as covariates for all ‘macaque only’ and ‘shared’ viruses. Error bars indicate the standard deviation between models initialized with different random seeds.

**Extended Data Fig. 9:**
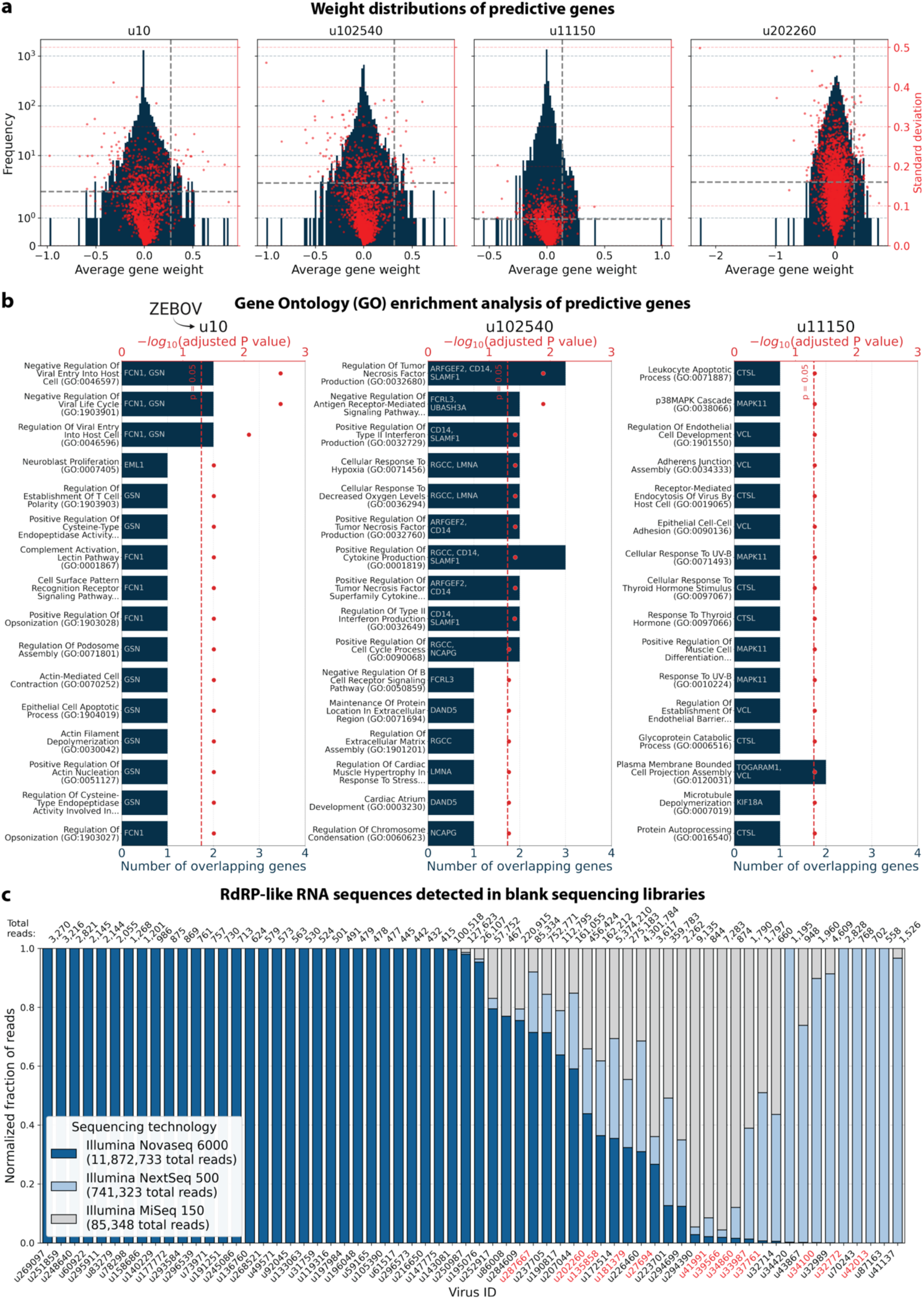
**a**, Average weight distributions of predictive genes from the models trained on highly variable genes with donor and time point as covariates for the four virus-like sequences with high predictive accuracy. The weights were averaged across models initialized using different random seeds and the standard deviations (SD) of the weights between seeds are shown in red. Gene weights were max normalized between random seeds before computing the average and SD. The dashed, grey lines indicate the minimum average gene weight and maximum SD for genes included in the enrichment analysis. **b**, Enrichment analysis of predictive genes from the regression model trained on highly variable genes with donor and time point as covariates. Approximately one third of the macaque Ensembl IDs did not have annotated gene names, which is a common problem for genomes from non-model organisms. We used gget^45^ to translate annotated Ensembl IDs to gene symbols and to perform enrichment analysis using Enrichr^71^ against the 2023 Gene Ontology (GO) Biological Processes database (‘GO_Biological_Process_2023)^72^. Gene names are listed on the bar plot. Reported P values were corrected with the Benjamini-Hochberg method. **c**, Sequencing reads were obtained by sequencing multiple ‘blank’ sequencing libraries containing only sterile water and reagent mix. The plot shows the fraction of reads that map to different virus IDs for each sequencing technology. The fractions were normalized to the total number of reads obtained for each technology. The data was generated by Porter et al.^63^ and analyzed using kallisto translated search with PalmDB. Virus IDs also detected in the macaque dataset are marked in red.

**Extended Data Fig. 10:**
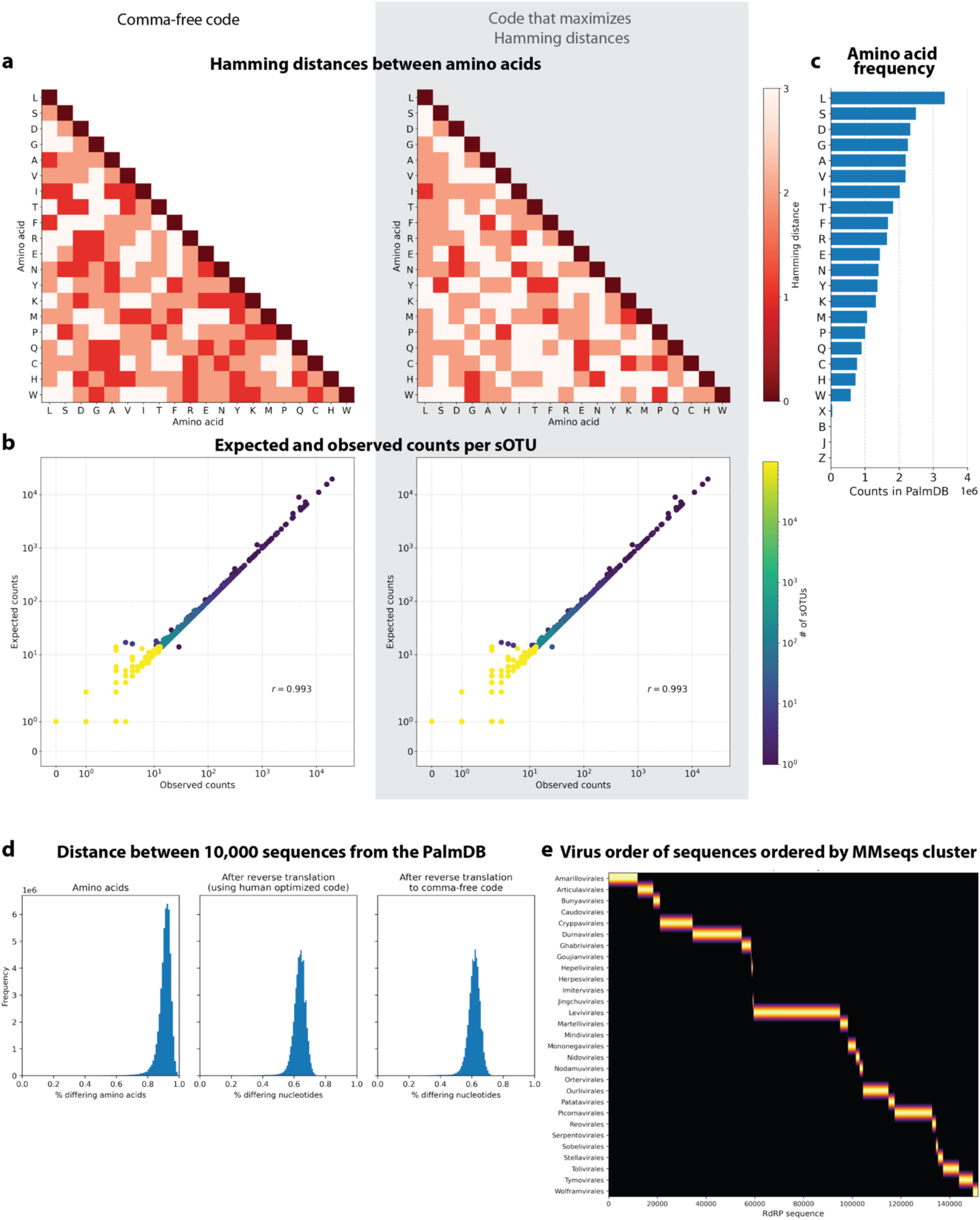
**a**, Hamming distances between amino acids in the comma-free code (left) and a second code that maximizes Hamming distances between amino acids that occur most often (right). **b**, We reverse translated all amino acid sequences in the PalmDB using the ‘standard’ genetic code (see Methods). The reverse translated PalmDB RdRP sequences were subsequently aligned to the optimized PalmDB amino acid reference (see Methods) with kallisto translated search. The left plot shows the expected and observed counts for each sOTU when kallisto performs the pseudoalignment in the comma-free code space. The plot on the right shows the expected and observed counts for each sOTU when kallisto performs the pseudoalignment using a second code that maximizes the Hamming distances between reverse translated amino acids. **c**, Occurrence of each amino acid in the PalmDB. **d**, Percentage of differing amino acids or nucleotides between 10,000 sequences randomly selected from the PalmDB before and after reverse translation using the standard genetic code (optimized for human) and comma-free code. **e**, The virus orders of RdRP sequences sorted based on their clustering by MMseqs2^79^ (see Methods).

